# Exploring the PET in vivo generator ^134^Ce as a theranostic match for ^225^Ac

**DOI:** 10.1101/2024.04.25.591165

**Authors:** David Bauer, Roberto De Gregorio, Edwin C. Pratt, Abram Bell, Alexa Michel, Jason S. Lewis

## Abstract

**Purpose:** The radionuclide pair cerium-134/lanthanum-134 (^134^Ce/^134^La) was recently proposed as a suitable diagnostic counterpart for the therapeutic alpha-emitter actinium-225 (^225^Ac). The unique properties of ^134^Ce offer perspectives for developing innovative in vivo investigations not possible with ^225^Ac. In this work, ^225^Ac- and ^134^Ce-labeled tracers were directly compared using internalizing and slow-internalizing cancer models to evaluate their in vivo comparability, progeny meandering, and potential as a matched theranostic pair for clinical translation. Despite being an excellent chemical match, ^134^Ce/^134^La has limitations to the setting of quantitative positron emission tomography imaging.

**Methods:** The precursor PSMA-617 and a macropa-based tetrazine-conjugate (mcp-PEG_8_-Tz) were radiolabelled with ^225^Ac or ^134^Ce and compared in vitro and in vivo using standard (radio)chemical methods. Employing biodistribution studies and positron emission tomography (PET) imaging in athymic nude mice, the radiolabelled PSMA-617 tracers were evaluated in a PC3/PIP (PC3 engineered to express a high level of prostate-specific membrane antigen) prostate cancer mouse model. The ^225^Ac and ^134^Ce-labeled mcp-PEG_8_-Tz were investigated in a BxPC-3 pancreatic tumour model harnessing the pretargeting strategy based on a trans-cyclooctene-modified 5B1 monoclonal antibody.

**Results:** In vitro and in vivo studies with both ^225^Ac and ^134^Ce-labelled tracers led to comparable results, confirming the matching pharmacokinetics of this theranostic pair. However, PET imaging of the ^134^Ce-labelled precursors indicated that quantification is highly dependent on tracer internalization due to the redistribution of ^134^Ce’s PET-compatible daughter ^134^La. Consequently, radiotracers based on internalizing vectors like PSMA-617 are suited for this theranostic pair, while slow-internalizing ^225^Ac-labelled tracers are not quantitatively represented by ^134^Ce PET imaging.

**Conclusion:** When employing slow-internalizing vectors, ^134^Ce might not be an ideal match for ^225^Ac due to the underestimation of tumour uptake caused by the in vivo redistribution of ^134^La. However, this same characteristic makes it possible to estimate the redistribution of ^225^Ac’s progeny noninvasively. In future studies, this unique PET in vivo generator will further be harnessed to study tracer internalization, trafficking of receptors, and the progression of the tumour microenvironment.

**TOC Graphic:** 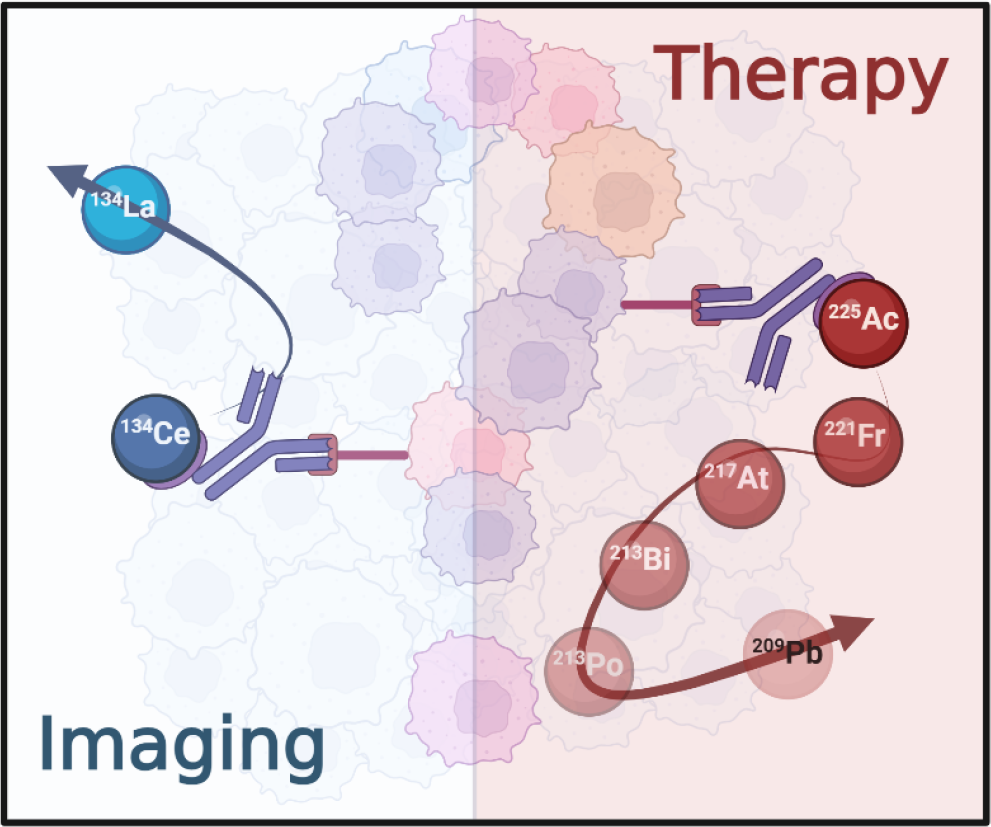

Redistribution of progeny. Investigating the ^225^Ac and ^134^Ce decay chain. This figure was created with BioRender.

## Introduction

Cerium-134 (^134^Ce, t_½_=3.2 d) is a fascinating new addition to the list of commercially available radionuclides. This imaging-compatible nuclide is currently in routine production at the U.S. Department of Energy Isotope Program to accelerate the development of theranostics pairs—matched radiopharmaceutical probes that serve as imaging and therapy agents [1]. ^134^Ce-based radiopharmaceuticals might be great candidates for clinical translation as positron emission tomography (PET) tracers [2, 3].

^134^Ce is not a typical PET radionuclide, since it lacks any utilizable radiation properties. It decays via electron capture to lanthanum-134 (^134^La, t_½_=6.5 min), and it is this short-lived daughter, ^134^La, that is the positron-emitter, decaying to stable barium-134 (Figure 1) [4]. For cancer diagnostics, ^134^Ce-based radiopharmaceuticals, with their relatively long physical half-lives, can deliver the radioactive payload to the targeted site. There, ^134^La, generated in vivo, allows for detection via PET imaging.

**Figure 1.**
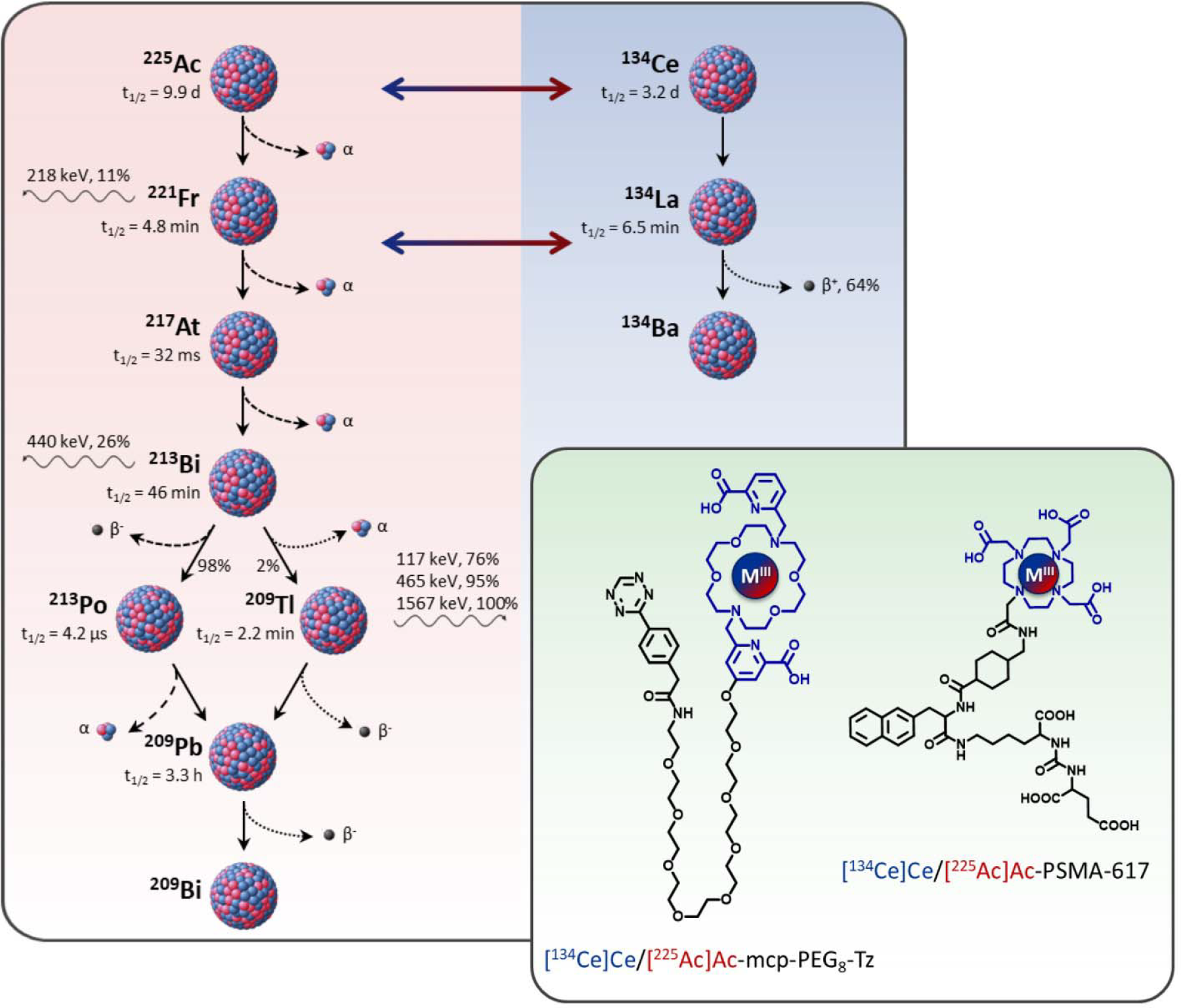
Decay scheme of ^225^Ac and ^134^Ce and structures of the investigated radioligands. While ^225^Ac and ^134^Ce show similar chemical behaviour and chelation chemistry, the PET-compatible progeny ^134^La can serve as an imaging surrogate for the similar short-lived daughter ^221^Fr.

This PET in vivo generator is currently undergoing preclinical investigation in cancer research as a diagnostic match for actinium-225 (^225^Ac, t_½_=9.9 d) [5-8]. Over the last decade, ^225^Ac has been developed into potent drug constructs targeting various forms of mostly highly aggressive and even treatment-resistant cancers. These constructs are currently being investigated in several clinical trials [9]. Despite its clinical success, one shortcoming of ^225^Ac-based therapies is that dosimetry is mainly extrapolated from one-point imaging data using an alternative radiotracer containing an imaging surrogate like gallium-68 or zirconium-89. From many perspectives, however, the PET in vivo generator ^134^Ce represents a superior theranostic match. The similar size and chemistry of actinium and cerium allow for the radiolabelling of the same precursors and the production of radiotracers with very similar pharmacokinetics. Additionally, the relatively long half-life of ^134^Ce allows for multiple imaging time points — ideal for preclinical studies — and also facilitates the synthesis and distribution of ^134^Ce-based imaging probes [10].

While ^134^Ce and ^225^Ac are undoubtedly a good chemical match, they may still be limited as imaging counterparts. Upon ^134^Ce’s decay, ^134^La is eliminated from the radiopharmaceutical due to the additional emission of Auger and inner conversion electrons, leaving the PET nuclide in a highly oxidated state, and eventually destroying the radiotracer [7]. Uncontained ^134^La might not be retained in the tumour environment, which could significantly affect image quantification.

In this study, ^134^La’s in vivo redistribution of two ^134^Ce-labeled radiotracers was evaluated (Figure 1). Additionally, the pharmacokinetics between the ^134^Ce- and ^225^Ac-labelled radiotracers were compared. Here, we demonstrate that quantitative imaging with ^134^Ce/^134^La is best applied to quickly internalizing radiotracers like PSMA-617 and is optimal for later imaging time points. However, when using slow-internalizing targeting strategies, e.g., with monoclonal antibodies (mAb), the image quantification is considerably impacted — even for later time points.

## Materials and Methods

All animal procedures were approved by the Institutional Animal Care and Use Committee (IACUC). [^134^Ce]CeCl_3_ and ^229^Th-derived [^225^Ac]Ac(NO_3_)_3_ were produced and supplied by the Department of Energy isotope (DOE) program. Additional information on the used instruments, synthesis of mcp-PEG_8_-Tz, analysis, in vitro and cell studies, and animal models can be found in the Supporting Information. The 5B1-TCO antibody conjugate (approximately three TCO moieties per mAb) was synthesized as previously published [11].

### Radiolabelling

The mcp-PEG_8_-Tz precursor was dissolved in dimethyl sulfoxide (DMSO) to obtain a stock solution of 10_– 3_ M (1 μL stock solution = 1 nmol precursor). A stock solution of 10–50 μL was added to 200 μL ammonium acetate buffer (0.25 M, pH = 5.5). Further, 0.19–1.1 MBq ^225^Ac or 18.5–111 MBq ^134^Ce, in the form of a stock solution (0.1 M HCL), were added. The reaction mixture was placed in a thermomixer (400 rpm) at 37.0 °C for 5 min.

PSMA-617 (5–20 μg) was radiolabelled similarly but incubated at 95°C for 10 min. Afterwards, the reaction was loaded on a C18 cartridge (Strata-X 33 μm Polymeric Reversed Phase, 30 mg/1 mL tubes), washed with PBS (1 mL), and eluted with ethanol (200 proof, 95%) to obtain the purified radiotracer.

The radiochemical conversion was determined via instant thin-layer chromatography (iTLC) using glass microfiber chromatography paper impregnated with silica gel (iTLC-SG, Agilent Technologies) and aqueous EDTA (ethylenediaminetetraacetic acid, 50 mM, pH 5.5) as the mobile phase. The iTLCs were scanned on a Bioscan AR-2000 radioTLC plate reader and analyzed using Winscan radio-TLC software (Bioscan Inc.). Radiochemical conversions > 95% were determined for all reactions. All ^225^Ac-based reactions were evaluated in equilibrium (> 8 h post-synthesis).

Serum stability studies in human serum at 37°C indicated that both radiotracers retain the radionuclide (> 95%) throughout 5 d (determined via radio iTLC).

### General information for the *in vivo* studies

All in vivo studies were performed in athymic nude mice. For pretargeting, female BxPC-3 tumour-bearing mice (tumour size 150–300 mm^3^) were intravenously injected (tail vain) with the trans-cyclooctene modified 5B1 (5B1-TCO) construct (100 μg in sterile filtered phosphate-buffered saline). After a three-day interval, radiolabelled mcp-PEG_8_-Tz (2 nmol per mouse) was intravenously administered (37 kBq for ^225^Ac, 3.7 MBq for ^134^Ce).

To evaluate the PSMA tracers, male PC3/PIP tumour-bearing mice (tumour size 150–300 mm^3^) were intravenously injected with radiolabelled PSMA-617 (0.5 μg per mouse) using the same activity levels.

For biodistribution studies, the mice were euthanized (CO_2_ asphyxiation, followed by cervical dislocation), and the tissues of interest were harvested and analysed on a Perkin Elmer Wizard^2^ 2480 Automatic Gamma Counter (Waltham, MA). ^225^Ac and ^134^Ce samples were measured for 2 min each, while ^213^Bi samples were measured for 30 s. ^225^Ac and ^213^Bi counts were both collected in the ^213^Bi-specific window (440 ± 80 keV) — the ^213^Bi data was collected directly after the biodistribution and the ^225^Ac data in equilibrium. The counts from each sample were corrected for decay and background and converted to %ID/g by comparison against internal standards. No decay correction was applied for ^213^Bi samples due to the complex ingrowth/decay relations (Supplemental Figure 1). The tissue collection and measurement of ^213^Bi in all selected samples was performed within 7 min after sacrificing the animal leading to a systematic error of less than 10%.

The data was analysed with the GraphPad Prism software 9.0 and represented as mean value ± standard error of the mean. The sample sizes were selected based on statistical considerations, ethical guidelines, and funding exigencies.

### Imaging

The mice were anaesthetized using 2% isoflurane for imaging. Positron emission tomography-computed tomography (PET-CT) images with ^134^Ce (approx. 3.7 MBq per mouse) were obtained on an Inveon™ PET-CT (Siemens) rodent scanner. For postmortem scans, the mice were euthanized (CO_2_ asphyxiation, followed by cervical dislocation) directly after completing the antemortem scan and re-imaged after a waiting period of 60 min (> nine half-lives of ^134^La, Supplemental Figure 2). All PET-CT images were analysed using the Inveon™ software suite. The counting rates in the reconstructed images were converted to the mean per cent injected dose per gram tissue (%ID/g) by applying a system-specific calibration factor (assuming 1 mL=1 g).

A Benchmark Jaszczak phantom was prepared with 336 μCi [^134^Ce]CeCl_3_ in 20 mL water containing 0.5% EDTA. It was imaged on the Inveon microPET system for 60 min, histogrammed, and reconstructed with a 512×512 pixel matrix after 12 iterations.

### Dosimetry estimates

The same dosimetry estimation method was used for both imaging and biodistribution data. Values of %ID/g were integrated using the trapezoidal method over each time point (1, 4, 24, 48, 72 h) with respect to seconds, yielding units of 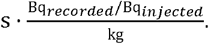 Activity after the final data point was extrapolated to infinity as an exponential decay function. An experimental internal radiation dose constant for each isotope of interest, given in energy output per Becquerel second 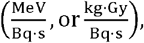, was taken from the National Nuclear Data Center [12]. This constant was multiplied by the integrated value to obtain the desired units of 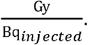 Thus, for any given measured injection, total administered dose in any organ of interest could be calculated.

## Results

### Biodistribution of free ^134^Ce

Before evaluating the in vivo behaviour of ^134^Ce- and ^225^Ac-based radiotracers, it was interesting to study the in vivo behaviour of unbound ^134^Ce. Therefore, 3.7 MBq [^134^Ce]CeCl_3_ was administered to healthy female nude mice. The mice were imaged at 4 and 24 time points, followed by terminal biodistribution (Figure 2). Both datasets show that free ^134^Ce predominantly accumulates in the liver and spleen with no substantial uptake in other tissues. A liver uptake of roughly 50 %ID/g was also reported for unchelated ^225^Ac [13]. However, Ce’s spleen uptake is approximately ten times higher than that observed for Ac. A potential explanation could be connected to cerium’s reversible redox chemistry, which allows it to change between trivalent and tetravalent forms [14].

**Figure 2.**
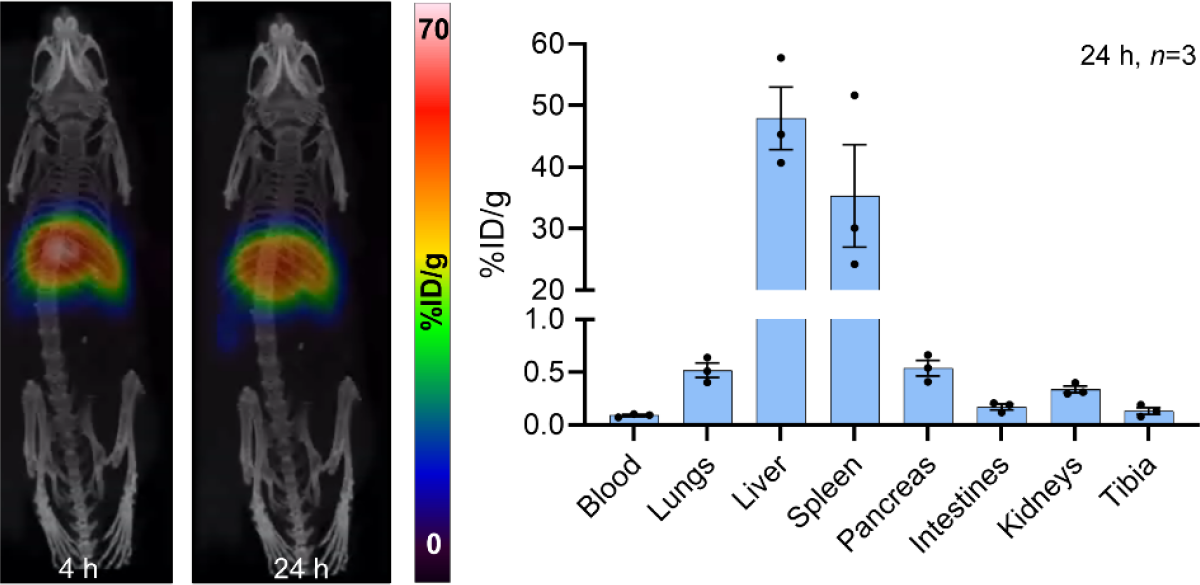
Biodistribution of [^134^Ce]CeCl_3_. Unbound ^134^Ce accumulates predominately in the liver and spleen as demonstrated (left) via PET imaging (maximum intensity projections at 4 and 24 h p.i.) and (right) terminal biodistribution (3.7 MBq per mouse).

However, the chemical environment — particularly a chelator — can stabilize the oxidation state of cerium. Hence, chelators with a very restricted coordination environment, like DOTA, DTPA and macropa, are reported to stabilize Ce(III), which then mimics the chemical behaviour of Ac(III), while chelators like 3,4,3-LI(1,2-HOPO) stabilize Ce(IV). In this nature, Ce(IV) has more chemical similarity to tetravalent metals like Th(IV) [5, 8, 10]. Hence, our studies focus on DOTA and macropa-based radiotracers.

### ^225^Ac and ^134^Ce-labeld PSMA-617

The DOTA-based radiotracer pair, ^225^Ac- and ^134^Ce-labelled PSMA-617 (Figure 3A), was evaluated in a PC3/PIP tumour mouse model for its in vivo behaviour. Since both radionuclides have an interesting decay chain, we additionally investigated the redistribution of their progeny. Figure 3B provides information about the location of [^225^Ac]Ac-PSMA-617 and its daughter ^213^Bi after injecting 37 kBq [^225^Ac]Ac-PSMA-617 in PC3/PIP-tumour bearing mice. ^213^Bi, with a half-life of 46 min, is the only alpha-emitter in ^225^Ac’s decay chain that can reasonably be quantified ex vivo. The data set indicates that unchelated ^213^Bi is mainly redistributed to the kidneys. Within a 3 d-interval, the progeny release progressively decreases, which can be ascribed to membrane turnover and tracer internalization. Figure 3C visualizes this trend. The substantial difference at the one-hour time point is partially a result of a certain amount of unchelated ^213^Bi present in the final formulation. At 1 h post-injection, 40% of the initial, free ^213^Bi remained in the subjects, mainly to be found in the kidneys [15]. Four hours post-injection, ^213^Bi is practically in equilibrium with ^225^Ac.

**Figure 3.**
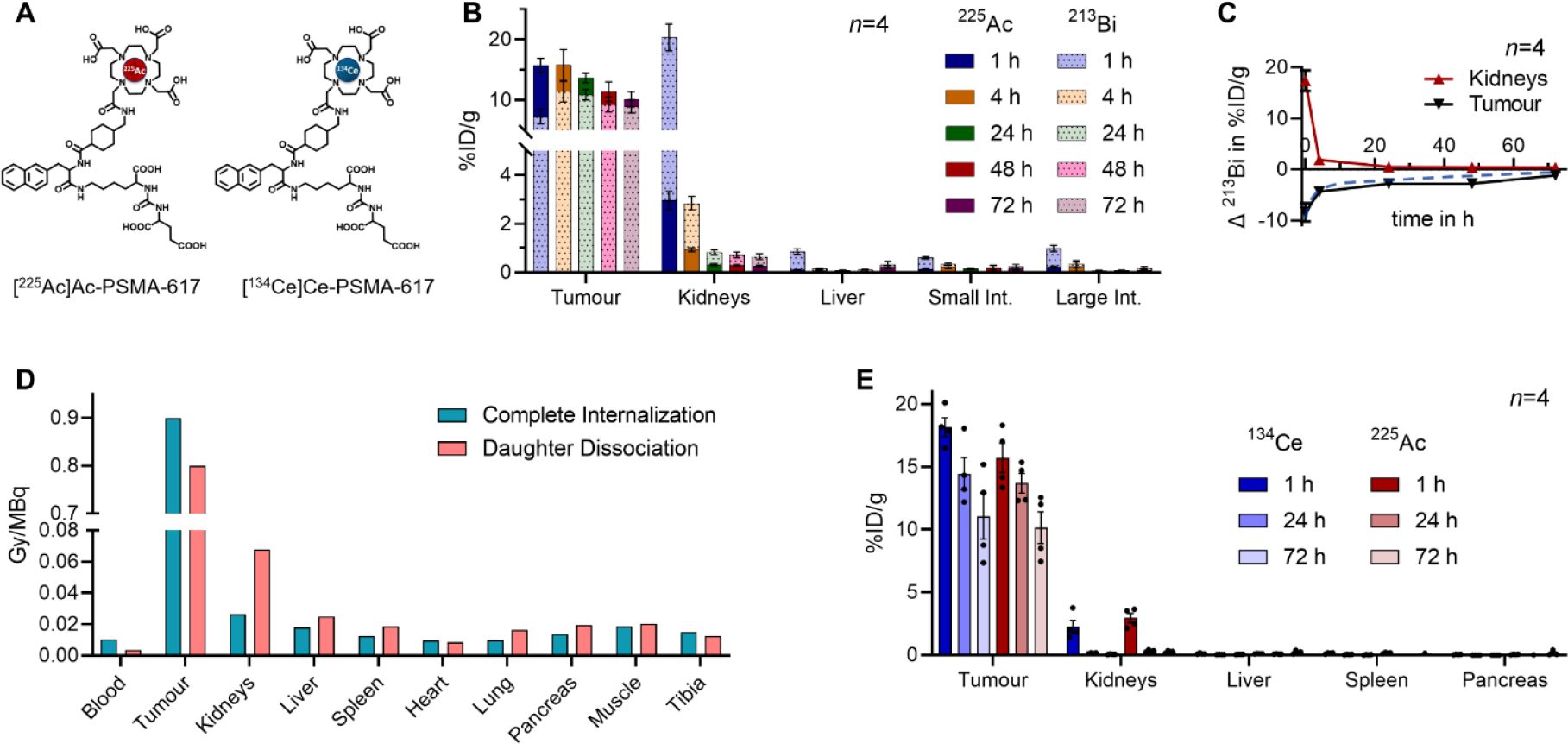
Investigation of PSMA-617 in a subcutaneous PC3/PIP tumour mouse model. **(A)** The theranostic pair [^225^Ac]Ac-PSMA-617 and [^134^Ce]Ce-PSMA-617. **(B)** Biodistribution data of ^225^Ac and ^213^Bi at selected time points after administration of 37 kBq [^225^Ac]Ac-PSMA-617 in the tissue of interest. **(C)** Plot of the relative ^213^Bi redistribution from tumour to kidneys in %ID/g. **(D)** Estimated relative biological effectiveness–weighted absorbed dose coefficients for [^225^Ac]Ac-PSMA-617 (in Gy-equivalent per MBq administered), disregarding and including the found redistribution of the progeny. **(E)** Direct comparison of [^225^Ac]Ac-PSMA-617 and [^134^Ce]Ce-PSMA-617 in the tissue of interest.

It is thus worth asking whether unchelated ^213^Bi should be seen as a risk factor and therefore removed before administering the dose. To answer this, note that 37 kBq of ^225^Ac equals a total of 1.8×10^11^ alpha emissions. Even in secular equilibrium, the same activity of ^213^Bi accounts for 1.5×10^8^ decays (0.08% of total alpha emissions). Hence, the initial level of unchelated progeny does not significantly impact the dosimetry. On the other hand, long-term redistribution of the daughter nuclides can do just that, as shown by the dosimetry estimates provided in Figure 3D. Biodistribution data (Figure 3 B) indicate increased kidney doses and reduced payloads at the tumour site. Figure 3E confirms that the biodistribution of ^225^Ac- and ^134^Ce-labelled PSMA-617 are virtually identical. However, PET imaging does not provide information about the location of [^134^Ce]Ce-PSMA-617 but exclusively about the daughter nuclide ^134^La, which is not retained in the radiopharmaceutical [7].

Finally, does [^134^Ce]Ce-PSMA-617 meet the expectations as a diagnostic partner? Figure 4A shows the maximum-intensity projections at 1, 24, and 48 hours after the injection of 3.7 MBq [^134^Ce]Ce-PSMA-617. At the 1 h timepoint, tumour uptake in the live mice (antemortem, AM) was determined to be 7 ± 2 %ID/g (*n*=4) via region-of-interest (ROI) analysis. This contradicts the biodistribution results (Figure 3E), indicating a tumour uptake of 18 ± 1 %ID/g. After each imaging time point, the cohort of mice was sacrificed and reimaged after 1 h to await the equilibrium between ^134^Ce and ^134^La. The analysis of the 1 h postmortem (PM) image indicates a tumour uptake of 16 ± 3 %ID/g, which affirms the biodistribution data for ^225^Ac- and ^134^Ce-labelled PSMA-617.

**Figure 4.**
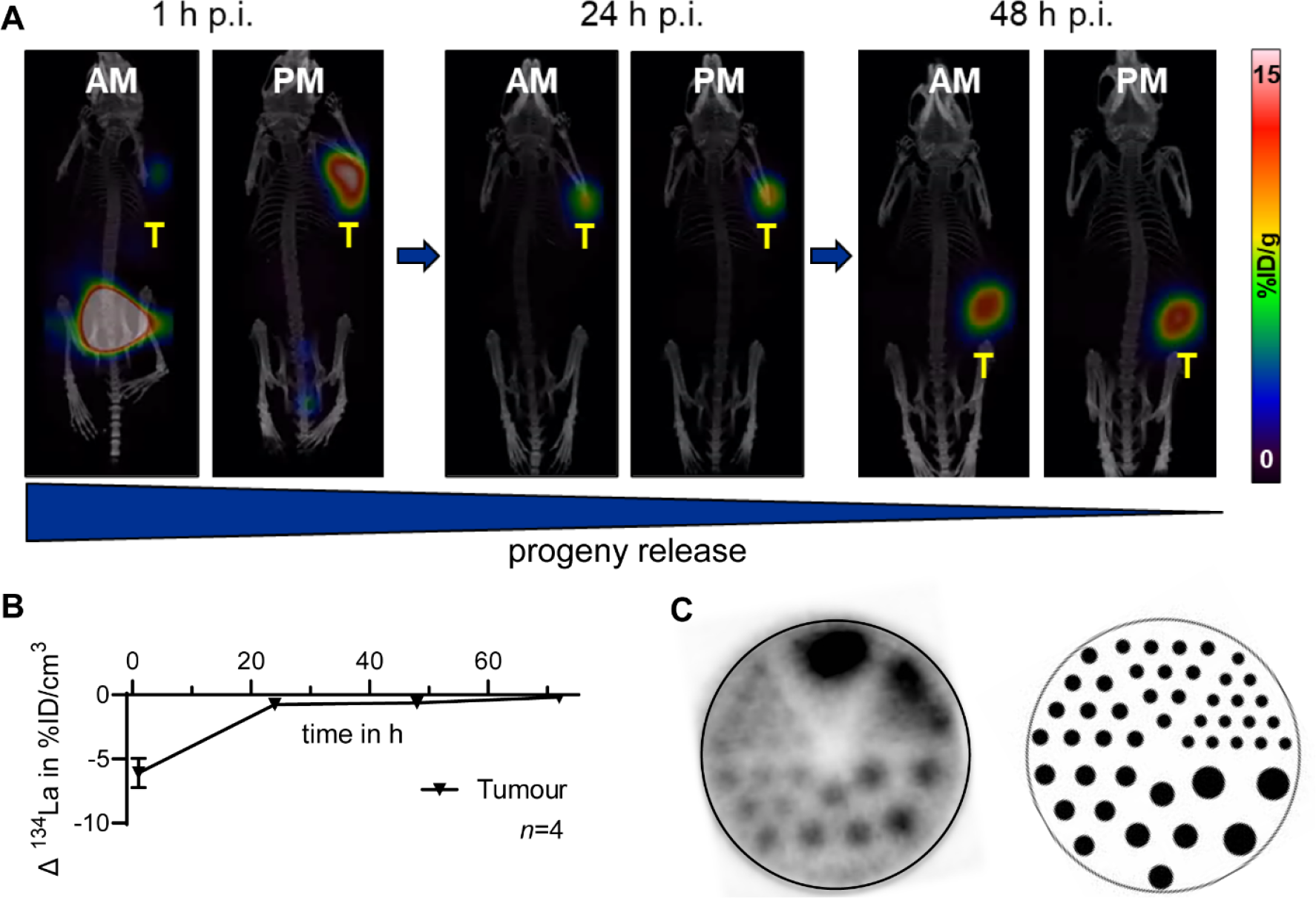
Evaluation of [^134^Ce]Ce-PSMA-617 via PET imaging of ^134^La. **(A)** PET imaging (maximum intensity projections) in a subcutaneous PC3/PIP tumour mouse model (3.7 MBq per mouse). Direct comparison of AM and PM imaging to visualize the redistribution of ^134^La at different time points. One out of four mice is represented. **(B)** Plot of the relative ^134^La redistribution from the tumours determined via comparative region-of-interest analysis of AM and PM images. **(C)** Maximum-intensity projection of a benchmark Jaszczak phantom (12.5 MBq [^134^Ce]CeCl_3_ in 20 mL).

This result implies that ^134^La, even though its half-life is just 6.5 min, redistributes significantly in the immediate post-injection hours. Upon membrane turnover and tracer internalization, the progeny release decreases, as indicated in Figure 4B. Consequently, PET imaging with ^134^Ce/^134^La does not perfectly reflect the biodistribution of ^225^Ac but gives a very conservative estimate that includes the potential redistribution of ^225^Ac’s daughters. At later time points, e.g. 48 h, PET imaging of [^134^Ce]Ce-PSMA-617 fairly reflects [^225^Ac]Ac-PSMA-617, mostly due to the internalization of the radiotracer.

### Jaszczak phantom

The ^134^Ce Jaszczak phantom shown in Figure 4C visualizes ^134^La’s emission of positrons with relatively high energy (E_mean_=1.2 MeV, E_max_=2.7 MeV), which results in a continuous slowing down approximation (CSDA) range in the water of 5.3 mm [16]. It should be emphasized that, as long as a good tumour-to-background ratio can be achieved, the possibility of identifying small lesions is not impaired by this range. Still, the quantitative accuracy of small structures might be compromised in the absence of range corrections [16]. Hence, small lesions, like the murine tumours evaluated here, might be slightly underestimated. To circumnavigate this issue, ROI sizes in dosimetry analysis were reduced to mitigate errors in perceived activity.

### Pretargeting with ^225^Ac and ^134^Ce

We sought to characterize differences in progeny redistribution between relatively fast- and slow-internalizing tracers. The slow-internalizing monoclonal antibody (mAb) 5B1, which targets the carbohydrate cell surface antigen 19-9 (overexpressed, e.g., in pancreatic ductal adenocarcinoma), is an excellent vector for this purpose. However, the direct labelling of this mAb would not be ideal since the long circulation time and slow tumour accumulation would not allow for visualization at early time points. It is expected that a ^134^Ce-labelled mAb, circulating in the bloodstream, cannot be accurately detected, since released ^134^La will be redistributed to the liver and spleen and partially be renally excreted. A pretargeting approach with the trans-cyclooctene-modified 5B1 and the tetrazine precursor, as illustrated in Figure 5, decouples the long biological half-life of the mAb from the delivery of the radioactive payload and allows for a relatively rapid clearance [17].

**Figure 5.**
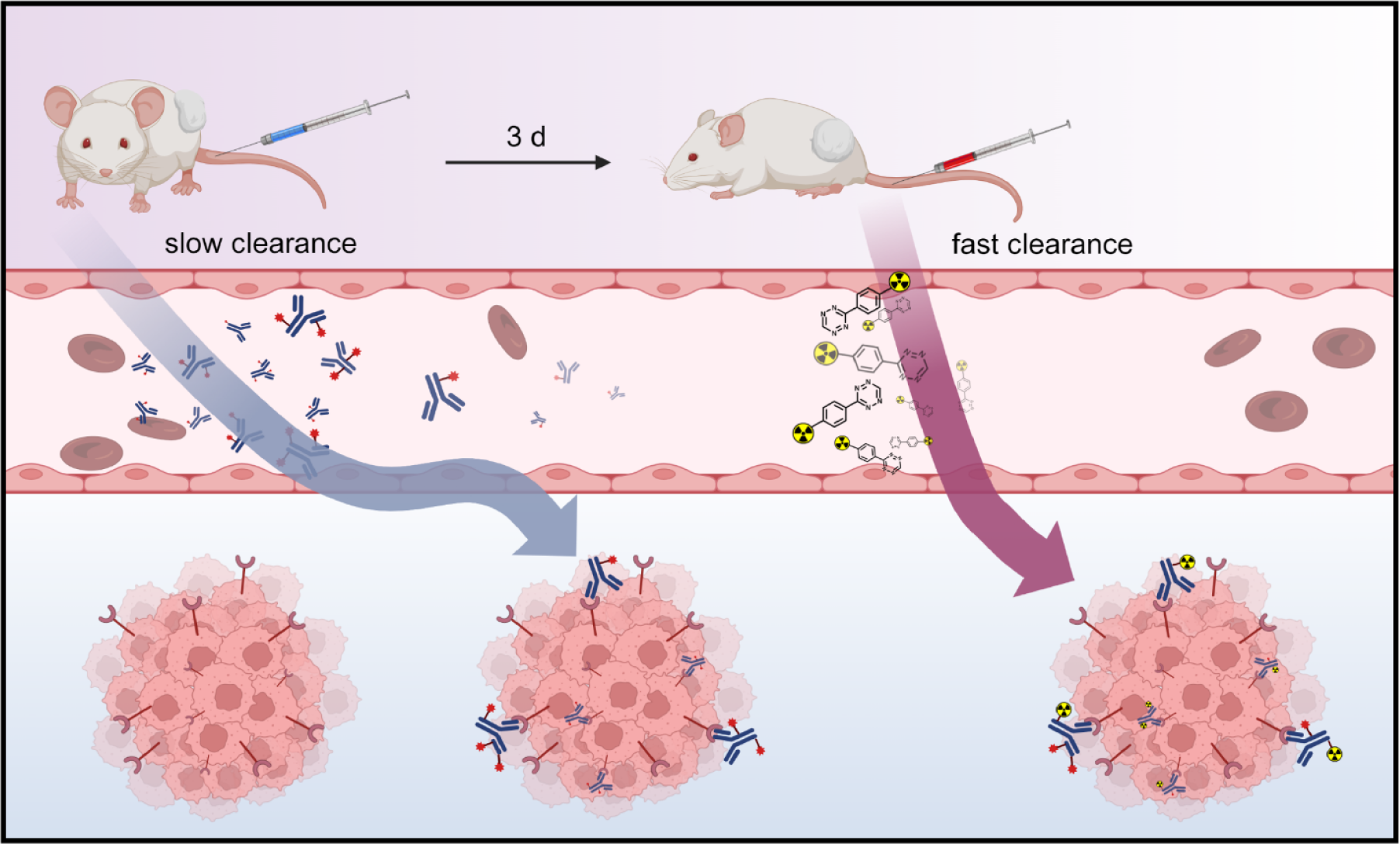
Illustration of the pretargeting approach, following the concept of reference [17]. The TCO (red star) modified mAb is administered, and after a certain time interval (here, 3 d), sufficient tumour uptake and clearance will be achieved. This is followed by injecting the radiolabelled tetrazine tracer — a small molecule that bio-orthogonally reacts with the TCO moiety. The tracer’s design allows for rapid renal clearance of unreacted excess. This figure was created with BioRender.

To facilitate this approach, the precursor mcp-PEG_8_-Tz was developed (Supplemental Scheme 1 and 2). As a chelator, macropa can be quantitatively radiolabelled within 5 min at room temperature for both ^225^Ac(III) and ^134^Ce(III), outcompeting DOTA by far.

The novel radiotracer, [^225^Ac]Ac-mcp-PEG_8_-Tz, was evaluated in a 5B1-TCO pretargeted BxPC-3 mouse model (Figure 6A) following earlier published procedures [15]. The biodistribution is comparable to previously investigated, similarly designed tetrazine [15, 18]. After 24 h, a maximal tumour uptake of 10.2 ± 0.8 %ID/g was determined. The blood values decrease relatively slowly (1.1 ± 0.4 %ID/g at 48 h), which can be attributed to a remaining amount of circulating 5B1-TCO clicking to the radiotracer within the bloodstream. Nevertheless, as recently explored, pretargeting enhances the onset of tumour uptake (here 4.2 ± 0.4 %ID/g at 1 h). Generally, it shows better background ratios than does direct labelling of the antibody [18, 19].

**Figure 6.**
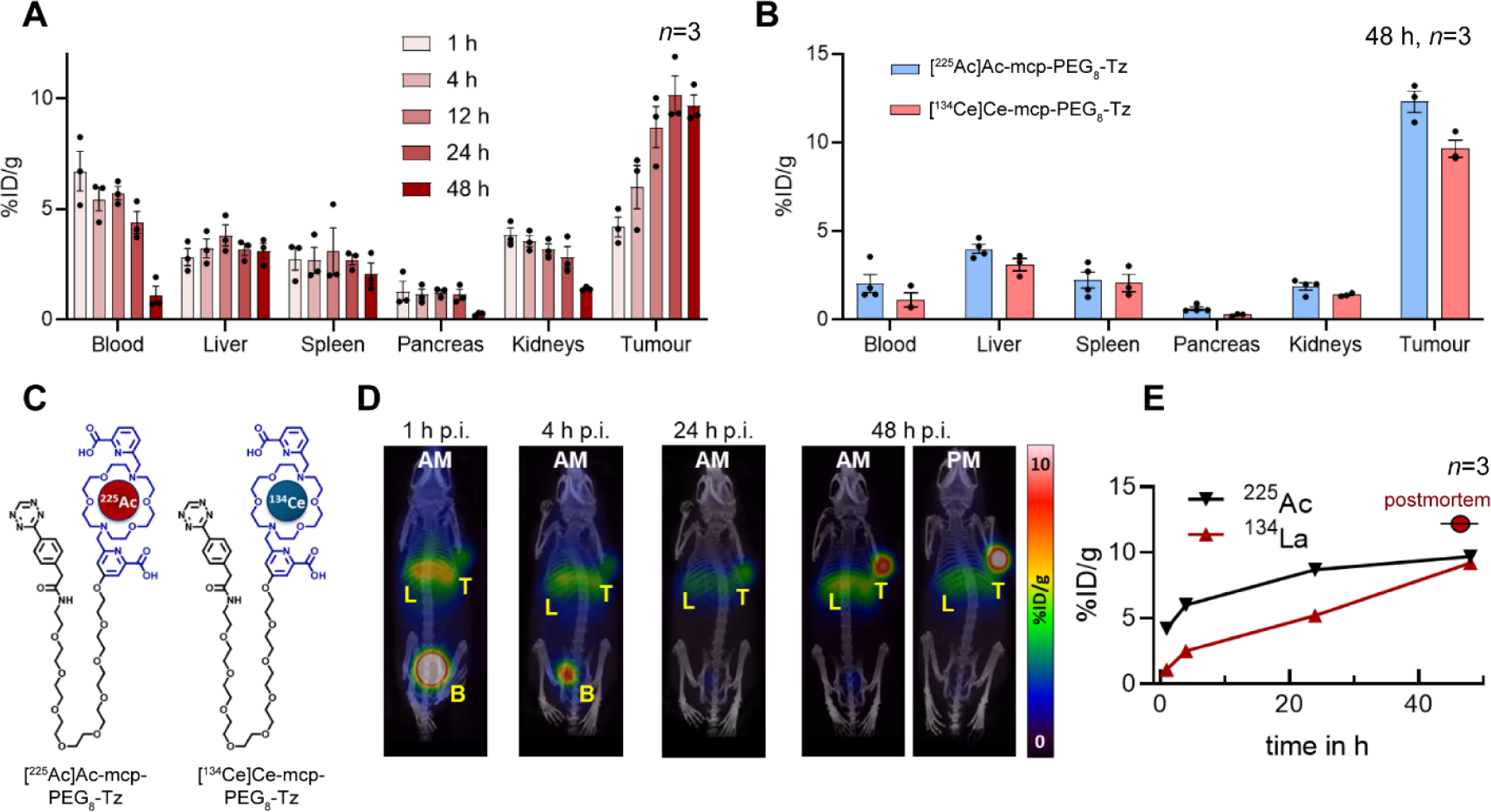
Investigating mcp-PEG_8_-Tz in a 5B1-TCO-pretargeted 5B1-TCO (100 μg, 0.7 nmol) subcutaneous BxPC-3 tumour mouse model. **(A)** Biodistribution data of [^225^Ac]Ac-mcp-PEG_8_-Tz (2 nmol, 37 kBq) in the tissue of interest. **(B)** Direct comparison of ^225^Ac- and ^134^Ce-labelled mcpPEG_8_-Tz via terminal biodistribution. **(C)** The theranostic pair [^225^Ac]Ac-mcp-PEG_8_-Tz and [^134^Ce]Ce-mcp-PEG_8_-Tz. **(D)** PET imaging (maximum intensity projections) in pretargeted mice receiving 3.7 MBq [^134^Ce]Ce-mcp-PEG_8_-Tz. Direct comparison of AM and PM imaging to visualize the redistribution of ^134^La at the terminal time point. **(E)** Direct comparison of the tumour uptake of ^225^Ac (via PM biodistribution, in equilibrium) and ^134^La (via AM ROI PET analysis) showing that PET imaging is underrepresenting the uptake, especially at early time points. The additional postmortem data point was determined via PM ROI PET analysis, highlighting the difference vs. the value-determined AM.

Together with the ^225^Ac cohort, four pretargeted mice were injected with [^134^Ce]Ce-mcp-PEG_8_-Tz. At the 48-h terminal time point, the theranostic pair was directly compared via biodistribution analysis (Figure 6B). With a tumour uptake of 12.3 ± 0.6 %ID/g at 48 h, the ^134^Ce-based tracer showed a slightly but significantly higher uptake than ^225^Ac, which reached a value of 9.7 ± 0.5 %ID/g (p-value=0.002, 2way ANOVA). No significant difference was observed in the other tissues.

Despite minor differences in their tumour uptake, ^225^Ac- and ^134^Ce-labelled Tz seem to function as a suitable theranostic pair. However, the ^134^La PET images (Figure 6D) make clear the impact of progeny redistribution. At the 24-hour timepoint, PET ROI analysis determined a tumour uptake of only 4.9 ± 0.5 %ID/g (AM), which increases to 9.1 ± 0.5 %ID/g (AM) at the 48 h timepoint. Mice were sacrificed and re-imaged PM. Without blood circulation, no redistribution of the daughter ^134^La occurred, and the tumour uptake value increased by 24%. In Figure 6E, the tumour uptake of [^225^Ac]Ac-mcp-PEG_8_-Tz, determined via biodistribution, is compared to the results from ^134^La PET imaging, showing a distinct underestimation of the delivered payload via PET imaging. The additional ‘equilibrium’ datapoint at 48 h marks the ^134^La/^134^Ce tumour uptake determined via imaging PM (11.3 ± 0.6 %ID/g), which agrees with the ex vivo biodistribution results (12.3 ± 0.6 %ID/g).

Over the course of 48 h, ^134^La’s redistribution considerably decreases but is still evident. The same must be true for ^225^Ac’s progeny. As in the PSMA-617 tracer study, ^213^Bi release was determined via ex vivo biodistribution at the terminal time point (48 h). While a tumour uptake of 9.7 ± 0.5 %ID/g was detected for [^225^Ac]Ac-mcp-PEG_8_-Tz, only 7.6 ± 0.3 %ID/g was determined for ^213^Bi (Supplemental Figure 3). To summarise, 22% of ^225^Ac’s progeny, ^213^Bi, and 19% of ^134^Ce’s daughter were redistributed in vivo at the 48-hour time point.

The reason for the steady decrease in daughter redistribution could be a combination of tracer internalization, membrane turnover, and encapsulation of the targeted cell into the tumour microenvironment. This can be partially verified with cell uptake studies. To test the tracer-internalization in vitro, the ‘pre-clicked’ mAb conjugate [^225^Ac]Ac-5B1-PEG_8_-mcp was incubated with 0.5×10^6^ BxPC-3 cells for 3 h either at 4 or at 37°C (Supplemental Figure 4). When the cells were incubated at 4°C with minimal cell metabolism, 83 ± 1% (*n*=3) of ^213^Bi was detected in the medium (not internalized), and only 65 ± 1% of ^213^Bi was detected when incubated at 37°C (Δ=22%).

## Discussion

This work demonstrates the relatively simple implementation of PET imaging with ^134^Ce/^134^La-based radiotracers and shows how this approach can add value to ^225^Ac-based studies. The high positron emission of 63%, ^134^Ce’s suitable half-life (t_½_=3.2 d), and its convenient coordination chemistry — compatible with standard chelators — are desirable properties for a clinical PET emitter. The relatively high positron energy could lead to quantitative underestimation of small lesions [16], but here, biodistribution data and PET ROI analysis (PM) were in reasonable accordance.

Unchelated [^134^Ce]CeCl_3_ and [^225^Ac]AcCl_3_ are mostly similar in their biodistribution in mice, with the major difference being the higher spleen uptake found for ^134^Ce. Once incorporated in a radiotracer, the radiometals make for a good pharmacokinetic match, as indicated in this study and recent literature [5, 6, 8, 10].

Radiolabelling of ^134^Ce is comparable to ^225^Ac, and radiopharmaceuticals containing the chelators DOTA or macropa are most suitable. However, due to its mild labelling conditions and fast kinetics, macropa should be considered the preferred choice.

This work also investigated whether the potential redistribution of the PET-compatible daughter, ^134^La, could impact the quantitative accuracy of ^134^Ce-based tracers. In general, fast-internalizing radiotracers (here, [^134^Ce]Ce-PSMA-617) are an acceptable theranostic match to their ^225^Ac twins, especially at later time points (24 h+), once internalization has progressed.

Interestingly, ^213^Bi’s release from [^225^Ac]Ac-PSMA-617-targeted tumours (Figure 3C) and ^134^La’s redistribution (Figure 4B) are quite comparable. However, ^213^Bi’s tumour release is slightly higher at all time points and, within 48 h, doesn’t decrease as fast as ^134^La’s. This could be related not only to ^213^Bi’s substantial longer half-life (t_½_=64 min), but also to the fact that ^213^Bi is part of a longer decay chain. Notably, ^225^Ac’s first daughter, francium-221 (^221^Fr, t_½_=4.8 min), has a similar half-life to ^134^La (t_½_=6.5 min). It can be hypothesized that since ^134^La does redistribute from the tumour tissue if not internalized, the same must be true for ^221^Fr. Currently, the common assumption is that due to ^221^Fr’s (and ^217^At’s) short half-life, the decay is located in situ with ^225^Ac. The in-vivo-generator ^134^Ce offers a new perspective on the redistribution of short-lived progeny.

Further, this pretargeting study using a novel tetrazine tracer illustrates that slowly internalizing radiotracers can present challenges when matching ^134^Ce with ^225^Ac, as the redistribution of ^134^La results in a significantly decreased PET tumour signal, even at later time points. However, in terms of clinical translation, it could be argued that PET imaging with ^134^Ce/^134^La will result in a quite conservative estimate of the therapeutic dose — which might not necessarily be a negative aspect. On the other hand, the toxicity profile of ^225^Ac-based tracer might not be accurately predicted with ^134^Ce since ^134^La seems to be quickly redistributed from its origin and partially accumulates in the liver (Supplemental Figure 5).

From a preclinical perspective, the possibility of AM/PM imaging with ^134^Ce-based tracers offers a convenient technique to investigate the internalization of tracers, including by estimating the delivered payload and radiotoxicity of ^225^Ac-based tracers (Figure 7).

**Figure 7.**
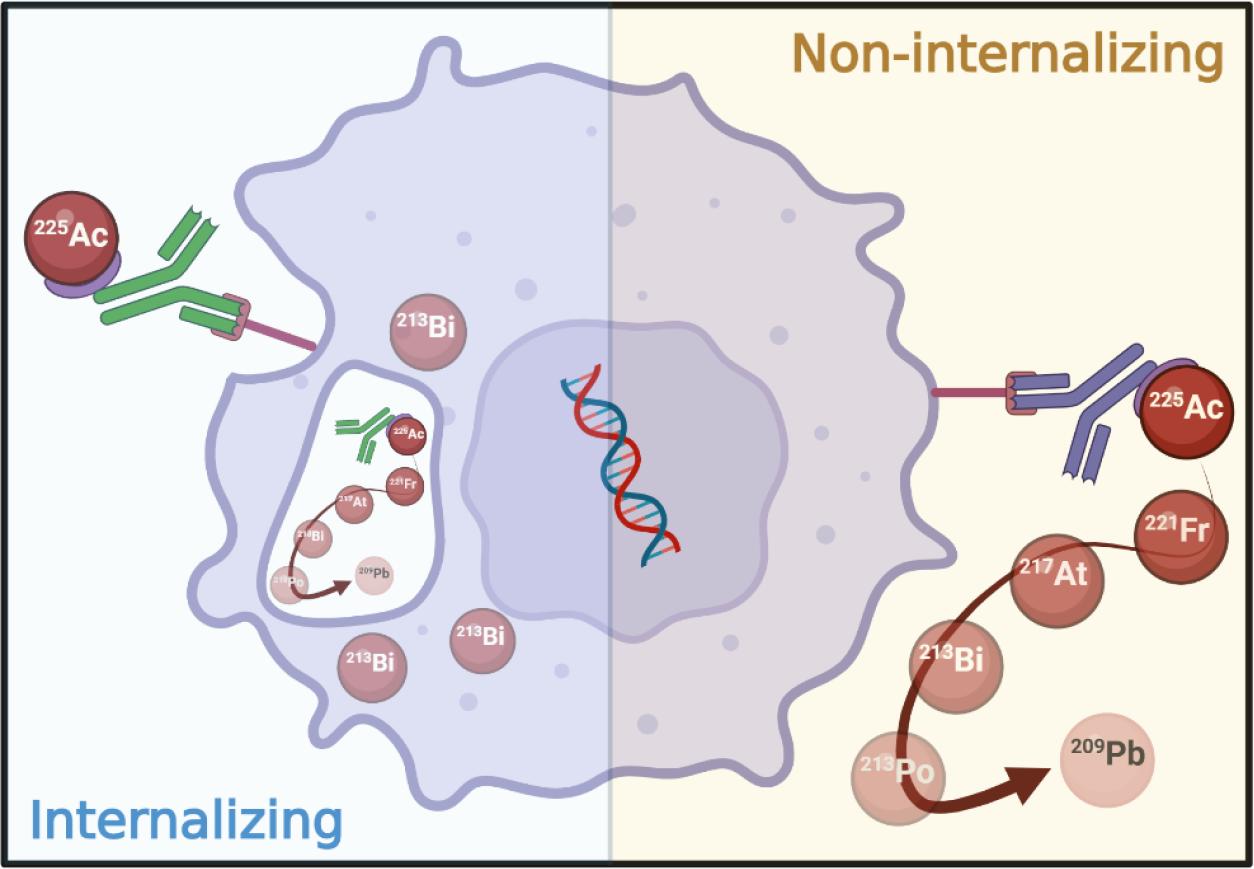
Impact on the progeny redistribution of internalizing versus non-internalizing radiotracers. This figure was created with BioRender.

## Conclusion

Whether or not ^225^Ac and ^134^Ce can serve as a theranostic match depends on the radiotracer employed. Ex vivo biodistribution comparisons are inadequate and require additional consideration to the progeny. The potential redistribution of the short-lived daughter, ^134^La, can be preclinically investigated by comparing AM and PM imaging. Overall, it is safe to assume that rapid internalizing radiotracers combined with imaging at later time points (+24 h) is the best strategy for this pair.

These findings also open exciting applications for PET in vivo generators, like ^134^Ce or ^140^Nd [7]. Due to ^134^La’s redistribution, ^134^Ce-based radiotracers can potentially facilitate the study of tracer internalization, trafficking of receptors, and the progression of the tumour microenvironment in vivo. Furthermore, since ^134^Ce and ^225^Ac can be considered a chemical match in an oncological setting, studying the redistribution of ^134^La (t_½_=6.5 min) from the tumour site might lead to an excellent predictor able to account for the redistribution of ^225^Ac’s first daughter, francium-221 (^221^Fr, t_½_=4.8 min), thereby improving dosimetry estimates.

## Declarations

### Ethics approval

All animal procedures were approved by the Institutional Animal Care and Use Committee (IACUC).

### Consent for publication

Not applicable.

### Availability of data and materials

The datasets generated during and/or analysed during the current study are available from the corresponding author upon reasonable request.

## Authors’ contributions

DB and JSL designed the research. DB performed the chemistry and in vitro experiments. DB, RDG, AB, and AM performed the in vivo experiments. RDG evaluated the nuclear magnetic resonance spectra. AB calculated the dosimetry estimates. ECP prepared and analysed the Jaszczak phantom. All authors joined in the writing of the manuscript and approved the final version.

## Acknowledgements

We thank the US Department of Energy Isotope Program (Isotope R&D Production) for providing the radionuclides. Furthermore, we thank George Sukenick and Rong Wang (MSK Analytical NMR Core Facility) for supporting NMR and mass spectrometry and Kishore Pillarsetty and Garon Scott (MSK Radiology) for their advice and support.

## References

1. National Isotope Development Center (November 17, 2022). Introducing a new isotope supply chain to the global market: Cerium-134. Accessed April 17, 2024. https://www.isotopes.gov/now_available_Cerium134.

2. Lubberink M, Lundqvist H, Tolmachev V. Production, PET performance and dosimetric considerations of 134Ce/134La, an Auger electron and positron-emitting generator for radionuclide therapy. Phys Med Biol. 2002;47:615–29. doi:10.1088/0031-9155/47/4/305.

3. Longtine M, Shim K, Abou D, Summer L, Voller T, Thorek D, Wahl R. Initial evaluation of Cerium-134 (134Ce) for immunoPET. Journal of Nuclear Medicine. 2022;63:2578.

4. Greenwood RC, Gehrke RJ, Helmer RG, Reich CW, Baker JD. Gamma-ray emission from 134Ce and levels in 134La. Nuclear Physics A. 1976;270:29–44. doi:10.1016/0375-9474(76)90125-1.

5. Bobba KN, Bidkar AP, Wadhwa A, Meher N, Drona S, Sorlin AM, et al. Development of CD46 targeted alpha theranostics in prostate cancer using 134Ce/225Ac-Macropa-PEG4-YS5. Theranostics. 2024;14:1344–60. doi:10.7150/thno.92742.

6. Bobba KN, Bidkar AP, Meher N, Fong C, Wadhwa A, Dhrona S, et al. Evaluation of 134Ce/134La as a PET Imaging Theranostic Pair for 225Ac alpha-Radiotherapeutics. J Nucl Med. 2023;64:1076–82. doi:10.2967/jnumed.122.265355.

7. Severin GW, Fonslet J, Kristensen LK, Nielsen CH, Jensen AI, Kjaer A, et al. PET in vivo generators 134Ce and 140Nd on an internalizing monoclonal antibody probe. Sci Rep. 2022;12:3863. doi:10.1038/s41598-022-07147-x.

8. Bailey TA, Wacker JN, An DD, Carter KP, Davis RC, Mocko V, et al. Evaluation of 134Ce as a PET imaging surrogate for antibody drug conjugates incorporating 225Ac. Nucl Med Biol. 2022;110-111:28–36. doi:10.1016/j.nucmedbio.2022.04.007.

9. Jang A, Kendi AT, Johnson GB, Halfdanarson TR, Sartor O. Targeted Alpha-Particle Therapy: A Review of Current Trials. Int J Mol Sci. 2023;24. doi:10.3390/ijms241411626.

10. Bailey TA, Mocko V, Shield KM, An DD, Akin AC, Birnbaum ER, et al. Developing the 134Ce and 134La pair as companion positron emission tomography diagnostic isotopes for 225Ac and 227Th radiotherapeutics. Nature Chemistry. 2020;13:284–9. doi:10.1038/s41557-020-00598-7.

11. Houghton JL, Zeglis BM, Abdel-Atti D, Sawada R, Scholz WW, Lewis JS. Pretargeted Immuno-PET of Pancreatic Cancer: Overcoming Circulating Antigen and Internalized Antibody to Reduce Radiation Doses. Journal of Nuclear Medicine. 2016;57:453–9. doi:10.2967/jnumed.115.163824.

12. National Nuclear Data Center (1994). Nuclear Decay Data in the MIRD Format. Accesed April 17, 2024. https://www.nndc.bnl.gov/nudat3/mird/

13. Yoshimoto M, Yoshii Y, Matsumoto H, Shinada M, Takahashi M, Igarashi C, et al. Evaluation of Aminopolycarboxylate Chelators for Whole-Body Clearance of Free 225Ac: A Feasibility Study to Reduce Unexpected Radiation Exposure during Targeted Alpha Therapy. Pharmaceutics. 2021;13. doi:10.3390/pharmaceutics13101706.

14. Piro NA, Robinson JR, Walsh PJ, Schelter EJ. The electrochemical behavior of cerium(III/IV) complexes: Thermodynamics, kinetics and applications in synthesis. Coordination Chemistry Reviews. 2014;260:21–36. doi:10.1016/j.ccr.2013.08.034.

15. Bauer D, Carter LM, Atmane MI, De Gregorio R, Michel A, Kaminsky S, et al. 212Pb-Pretargeted Theranostics for Pancreatic Cancer. J Nucl Med. 2024;65:109–16. doi:10.2967/jnumed.123.266388.

16. Carter LM, Kesner AL, Pratt EC, Sanders VA, Massicano AVF, Cutler CS, et al. The Impact of Positron Range on PET Resolution, Evaluated with Phantoms and PHITS Monte Carlo Simulations for Conventional and Non-conventional Radionuclides. Mol Imaging Biol. 2020;22:73–84. doi:10.1007/s11307-019-01337-2.

17. Altai M, Membreno R, Cook B, Tolmachev V, Zeglis BM. Pretargeted Imaging and Therapy. J Nucl Med. 2017;58:1553–9. doi:10.2967/jnumed.117.189944.

18. Keinanen O, Fung K, Brennan JM, Zia N, Harris M, van Dam E, et al. Harnessing 64Cu/67Cu for a theranostic approach to pretargeted radioimmunotherapy. Proc Natl Acad Sci U S A. 2020;117:28316–27. doi:10.1073/pnas.2009960117.

19. Poty S, Carter LM, Mandleywala K, Membreno R, Abdel-Atti D, Ragupathi A, et al. Leveraging Bioorthogonal Click Chemistry to Improve 225Ac-Radioimmunotherapy of Pancreatic Ductal Adenocarcinoma. Clin Cancer Res. 2019;25:868–80. doi:10.1158/1078-0432.CCR-18-1650.

