## Supporting Information for "Exploring the PET in vivo generator ^134^Ce as a theranostic match for ^225^Ac"

### 1. Additional materials and methods

#### 1.1 General laboratory equipment

All instruments were calibrated and maintained in accordance with standard quality-control procedures. A Capintec CRC-55tR Dose Calibrator (Capintec, Ramsey, NJ) was used for all activity measurements except for those of harvested tissues. Nuclear magnetic resonance (NMR) spectroscopy was performed on a Bruker Avance III UltraShield Plus 500 MHz NMR equipped with a BBO probe. All NMR spectra were measured in Dimethyl sulfoxide-$d_6$ and evaluated using the MestReNova software (Version: 6.0.2-5475). Liquid chromatography-mass spectrometer (LC-MS) was performed on a C18 BEH column (100 mm with 1.7 µm particle size) using a Waters ACQUITY UPLC (Milford, MA, USA) interfaced to a single quadrupole ESI (electrospray ionization) mass spectrometer (SQD), Evaporative Light Scattering Detector (ELSD), and Photo Diode Array (PDA) Detector. Antibody quantification was performed on a Thermo Scientific NanoDrop One Spectrophotometer.

#### 1.2 Cell line and animal model

The pancreatic cancer cell line BxPC-3 (American Type Culture Collection, ATCC) was cultured according to the recommended conditions at 5% $CO_2$ atmosphere and 37°C in RPMI-1640 medium containing 10% foetal bovine serum. The prostate cancer cell line PC3/PIP (PC3 engineered to express a high level of Prostate Specific Membrane Antigen, MSKCC) was cultured according to the recommended conditions at 5% $CO_2$ atmosphere and 37°C in RPMI-1640 medium containing 10% foetal bovine serum. The MSK Media Preparation Core provided the media. For the subcutaneous xenografts, the cells were stripped in the absence of magnesium or calcium ions using a mixture of 0.25% trypsin and 0.5 mM EDTA in Hank's Balanced Salt Solution and concentrated in 1 mL of the corresponding medium. A small aliquot was used to determine the cell count (Beckman Coulter Vi-CELL XR). For the xenografts, the remaining cells were diluted with the medium so that 50 µL contained approximately $3\times10^6$ cells for BxPC-3 and $1\times10^6$ for PC3/PIP.

The studies were performed in female (BxPC-3) and male (PC3/PIP) athymic nude mice (Crl:NU(NCr)-Foxn1$^{nu}$) obtained from Charles River Laboratories (Stone Ridge, NY). After arrival, the mice were kept in the MSK vivarium for 1 wk before any experimental handling was performed. The animals were allowed free access to water and food, and all animal care and experimental procedures were approved by the Institutional Animal Care and Use Committee (IACUC). Before the treatment, the mice were on a Sulfatrim diet to prevent skin infections. For the subcutaneous xenografts, a 1:1 ratio of Corning Matrigel Matrix and the cell solution was prepared and stored on ice for the injections. Each mouse received a subcutaneous injection of 100 µL (50 µL cell solution, $3\times10^6$ cells, viability > 95%, + 50 µL Matrigel) into the flank. The injections were performed under anaesthesia (2% isoflurane, Baxter Healthcare, Deerfield, IL, USA).

Palpable tumours of similar size (100–200 mm$^3$) developed 4–5 wk after grafting. The radiotracer and 5B1-TCO administration were performed via tail vein injection.

#### 1.3 Synthesis of mcp-PEG$_8$-Tz

All chemicals not further specified were purchased from Sigma-Aldrich and used as received. The identity and purity of the synthesized compounds were confirmed via $^1$H NMR, $^{13}$C NMR and ESI-MS.

The HPLC purifications (Solvent A: 0.1% trifluoroacetic acid (TFA) in water, Solvent B: 0.1% TFA in acetonitrile) were performed on a Shimadzu UFLC HPLC system equipped with a DGU-20A$_3$ degasser, an SPD-M20A UV detector, a LC-20AB pump system, and a CBM-20A communication BUS module, using a C18 reversed phase XTerra® Preparative MS OBD column (10 µm, 19×250 mm) at a constant flowrate of 8 mL/min.

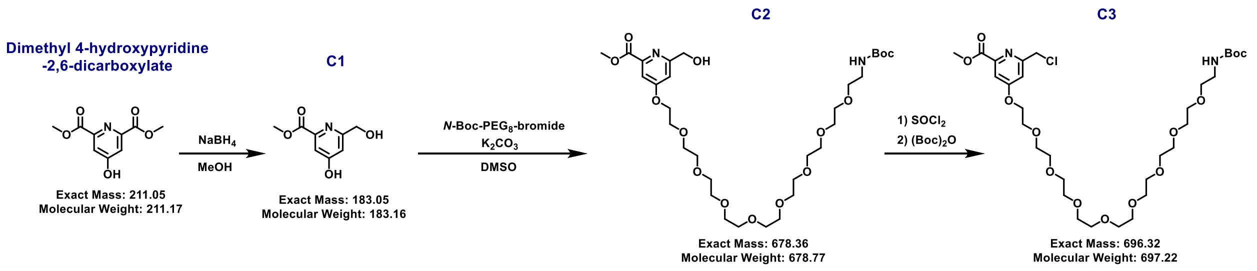

**Supplemental Scheme 1.** Synthesis overview for compound **C4**.

**C1:** Dimethyl 4-hydroxypyridine-2,6-dicarboxylate (Ambeed Inc., 2.0 g, 9.5 mmol) was suspended in anhydrous methanol (50 mL), and the flask was placed on ice. $NaBH_4$ (360 mg, 9.5 mmol) was added, and the conversion was monitored via TLC (thin-layer chromatography). More $NaBH_4$ was added stepwise over the course of 6 h till the starting material was mostly consumed. After the removal of the solvent, the crude product was purified via silica column chromatography (chloroform/methanol 0→15%).

**C1** was obtained as a colourless solid (435 mg, 25%). $^1$H NMR (500 MHz, DMSO-d$_6$): δ = 7.27 (d, $^4J$ = 2.0 Hz, 1H, H$_{Ar}$), 7.07 (d, $^4J$ = 2.0 Hz, 1H, H$_{Ar}$), 4.52 (s, 2H,CH$_2$), 3.85 (s, 3H,CH$_3$) ppm.$^{13}$C NMR (126 MHz, DMSO-d$_6$): δ =166.76 (C$_{Ar}$), 164.77 (C=O), 163.39 (C$_{Ar}$), 146.89 (C$_{Ar}$), 111.59 (CH$_{Ar}$), 110.85 (CH$_{Ar}$), 63.11 (CH$_2$), 52.48 (CH$_3$) ppm. TFA is visible in the spectrum (δ = 158 (q, $^3J$ = 34 Hz) and 116 (q, 296 Hz) ppm) since the NMR was measured with a HPLC purified fraction. ESI-MS — m/z calculated for $C_9H_9NO_5$: 211.0 u; found: [M+H]$^+$ 212.1.

**C2:** C1 (200 mg, 1.1 mmol) and N-Boc-PEG$_8$-bromide (BroadPharm, 630 mg, 1.1 mmol) were dissolved in anhydrous dimethyl sulfoxide (DMSO, 5 mL), and $K_2CO_3$ (75 mg, 0.55 mol) was added. The mixture was stirred at 70°C for 2 d (TLC control). After cooling to rt (room temperature), the reaction was filtered, and the filtrate was diluted with water (1:1). The mixture was directly injected into the HPLC (400 µL per run). The collected fractions were combined, frozen, and lyophilized.

**C2** was obtained as a colourless oil (391 mg, 36%, TFA salt). $^{1}H$ NMR (500 MHz, DMSO-$d_6$): δ = 7.44 (d, $^{4}J$ = 2.1 Hz, 1H, $H_{Ar}$), 7.23 (s, 1H, $H_{Ar}$), 6.74 (s, 1H,NHC=O), 4.56 (s, 2H, $CH_2$), 4.31–4.22 (m, 2H,$OCH_2$), 3.80–3.70 (m, 2H,$OCH_2$), 3.50 (s, 28H, $OCH_2$), 3.36 (m, 2H ,$OCH_2$), 3.09–3.01 (m, 2H, $CH_2$), 1.37 (s, 9H, $(CH_3)_3$) ppm. $^{13}C$ NMR (126 MHz, DMSO-$d_6$): δ = 166.10 ($C_{Ar}$), 165.14 (C=O), 164.44 ($C_{Ar}$), 155.57 (C=O), 148.23 ($C_{Ar}$), 109.82 ($CH_{Ar}$), 109.37 ($CH_{Ar}$), 77.57 ($\mathit{C}(CH_3)_3$), 69.70–69.85 (15×$OCH_2$), 69.16 ($OCH_2$), 68.52 ($OCH_2$), 67.79 ($CH_2$), 63.75 ($CH_2$), 52.41 ($CH_3$), 28.23 ($C(\mathit{C}H_3)_3$) ppm. ESI-MS — m/z calculated for $C_{31}H_{54}N_2O_{14}$: 678.4 u; found: $[M+H]^+$ 679.5.

**C3:** C2×TFA (200 mg, 0.25 mmol) was added to a flask and placed on ice before adding thionyl chloride (1 mL). After 30 min, excess of thionyl chloride was removed under vacuum. The obtained oil was a mixture of product (C3) and Boc-deprotected product. The oil was dissolved in dichloromethane (1 mL), and di-tert-butyl decarbonate (1 mL) was added. After 30 min, excess of the reactant was removed under vacuum. The residue was dissolved in water/acetonitrile (5 mL each) and injected into the HPLC (400 µL per run). The collected fractions were combined, frozen, and lyophilized.

**C3** was obtained as a colourless oil (148 mg, 73%, TFA salt). $^{1}H$ NMR (500 MHz, DMSO-$d_6$): δ = 7.53 (d, $^{4}J$ = 1.2 Hz, 1H, $H_{Ar}$), 7.39 (s, 1H, $H_{Ar}$), 6.74 (s, 1H, NHC=O), 4.76 (s, 2H, $CH_2$), 4.34–4.22 (m, 2H, $OCH_2$), 3.93–3.83 (m, 3H, $OCH_2$), 3.84–3.73 (m, 2H, $OCH_2$), 3.65–3.42 (m, 30H, $OCH_2$), 3.40–3.33 (m, 2H, $OCH_2$), 3.05 (d, 2H, $CH_2$), 1.37 (s, 9H, $(CH_3)_3$) ppm. $^{13}C$ NMR (126 MHz, DMSO-$d_6$): δ =166.21 ($C_{Ar}$), 164.86 (C=O), 158.39 ($C_{Ar}$), 155.57 (C=O), 149.03 ($C_{Ar}$), 112.98 ($CH_{Ar}$), 110.84 ($CH_{Ar}$), 77.56 ($\mathit{C}(CH_3)_3$), 70.06–69.38 (15×$OCH_2$), 69.15 ($OCH_2$), 68.48 ($CH_2$), 68.04 ($CH_2$), 52.53 ($CH_2$), 46.28 ($CH_2$), 28.22 ($C(\mathit{C}H_3)_3$) ppm. ESI-MS — m/z calculated for $C_{31}H_{53}ClN_2O_{13}$: 696.3 u; found: $[M+H]^+$ 697.4.

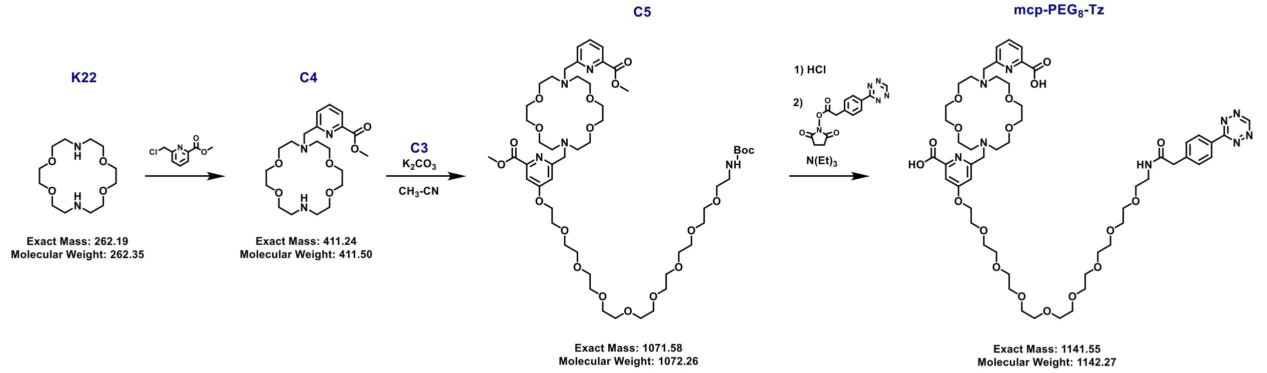

**Supplemental Scheme 2.** Synthesis overview for **mcp-PEG$_8$-Tz**.

**C4:** Kryptofix®22 (K22, 100 mg, 0.38 mmol) and methyl 6-(chloromethyl) picolinate (Chemscene, 85 mg, 0.46 mol) were dissolved in anhydrous DMSO (1 mL) and triethylamine (0.21 mL) was added. The mixture was stirred at 70°C overnight (TLC control). After cooling to rt, water was added (4 mL), and the solution was directly injected into the HPLC (800 µL per run). The collected fractions were combined, frozen, and lyophilized.

**C4** was obtained as a colourless solid (121 mg, 61%, TFA salt). The NMR and MS data was in accordance with previously published literature (F. Reissig, et al. Cancers. 2021;13(8):1974. doi:10.3390/cancers13081974).

**C5: C3**×TFA (50 mg, 37 µmol) and **C4**×TFA (29 mg, 55 µmol) were dissolved in acetonitrile (1 mL), and $K_2CO_3$ (31 mg, 0.22 mol) was added. The mixture was stirred at 70°C for 2 d (TLC control). After cooling to rt, the reaction was filtered, and the filtrate was diluted with water to a total of 5 mL. The mixture was directly injected into the HPLC (800 µL per run). The collected fractions were combined, frozen, and lyophilized.

**C5** was obtained as a colourless oil (9.6 mg, 22%, TFA salt). $^1$H NMR (500 MHz, DMSO-d$_6$): δ = 8.10 (s, 2H, 2×$CH_{Ar}$), 7.81 (d,1H, $CH_{Ar}$), 7.61 (s, 1H, $CH_{Ar}$), 7.43 (s, 1H, $CH_{Ar}$), 6.74 (s, 1H, NHC=O), 4.62 (d, 2H, $CH_2$), 4.59–4.42 (m, 2H, $CH_2$), 4.30 (s, 2H, $OCH_2$), 3.89 (m, 6H, 2×$CH_3$ ), 3.87–3.75 (m, 9H, $OCH_2$), 3.59 (s, 18H, $OCH_2$), 3.56–3.43 (m, 27H, $OCH_2$), 3.36 (t, 3H, $OCH_2$), 3.05 (m, 2H, $CH_2$), 1.36 (s, 9H, $(CH_3)_3$) ppm. $^{13}$C NMR (126 MHz, DMSO-d$_6$): δ = 166.25 ($C_{Ar}$), 164.64 (2×C=O), 155.58 (C=O), 148.99 ($C_{Ar}$), 147.13 ($C_{Ar}$), 139.08 ($CH_{Ar}$), 128.34 ($CH_{Ar}$), 124.89 ($CH_{Ar}$), 111.39 ($CH_{Ar}$), 77.57 ($C(CH_3)_3$), 70.14–68.10 (25×$OCH_2$), 64.59 (2× $CH_2N$), ) 56.92 (8×$CH_2$), 53.27 (2×$CH_3$), 52.71 ($CH_3$), 45.72 ($CH_2$), 28.22 ($C(CH_3)_3$) ppm. Signals for 1×$CH_{Ar}$ and 2×$C_{Ar}$ are not visible. TFA is visible in the spectrum (δ = 158 (q, $^3J$ = 34 Hz) and 117 (q, 296 Hz) ppm). ESI-MS — m/z calculated for $C_{51}H_{85}N_5O_{19}$: 1071.6 u; found: $[M+H]^+$ 1072.7, $[M+Na]^+$ 1093.8.

**mcp-PEG$_8$-Tz: C5**×TFA (8 mg, 6.7 µmol) was dissolved in HCl (1 mL, 0.1 M) and heated to 50°C for 3 h. The complete deprotection (2×methyl and Boc) was monitored via LC-MS. The mixture was frozen and lyophilized to obtain a colourless solid, which was dissolved in acetonitrile (1 mL). Tetrazine-NHS ester (Broad Pharm, 2.5 mg, 8.0 µmol) and triethylamine (5 µL) were added, and the reaction was reacted at rt for 3 h. Acidified water (0.1% TFA, 2 mL) was added (tetrazines are sensitive to basic conditions), and the solution was directly injected into the HPLC (800 µL per run). The collected fractions were combined, frozen, and lyophilized.

**Mcp-PEG$_8$-Tz** was obtained as a bright pink solid (3.7 mg, 44%, TFA salt). $^1$H NMR (500 MHz, DMSO-d$_6$): δ = 10.58 (s, 1H, $C_{Tz}H$), 8.43 (m, 2H, 2×$CH_{Ar}$), 8.25 (s, 1H, NHC=O), 8.18–8.06 (m, 2H, 2×$CH_{Ar}$), 7.78 (d, 1H, $CH_{Ar}$), 7.59 (d,1H, $CH_{Ar}$), 7.55 (d, 2H, 2×$CH_{Ar}$), 7.40 (s, 1H, $CH_{Ar}$), 4.67 (d, 2H, $CH_2$), 4.59 (s, 2H, $CH_2$), 4.31 (s, 2H, $CH_2$), 3.84 (s, 8H, $CH_2$), 3.79 (s, 2H, $OCH_2$), 3.57 (m,12H, $OCH_2$), 3.56–3.46 (m, 28H, $OCH_2$), 3.45–3.40 (m, 2H, $OCH_2$), 3.30–3.18 (m, 4H, $CH_2$) ppm. $^{13}$C NMR (126 MHz, DMSO-d$_6$): δ =169.55 ($C_{Ar}$), 166.49 ($C_{Ar}$), 165.46 (C=O), 165.34 – 165.24 (2×C=O), 147.78 ($C_{Ar}$), 141.78 ($C_{Ar}$), 139.16 ($C_{Ar}$), 130.12 (2×$CH_{Ar}$), 127.65 (2×$CH_{Ar}$), 124.51 ($CH_{Ar}$), 110.24 ($CH_{Ar}$), 70.98 – 67.77 (24×OCH2), 64.53 ($CH_2$), 57.04 – 56.24 (8×$NCH_2$), 53.39 ($CH_2$), 41.75 ($CH_2$) ppm. TFA is visible in the spectrum (δ = 158 (q, $^3J$ = 34 Hz) and 117 (q, 296 Hz) ppm). Not all carbon signals are visible in the spectrum. TFA is visible in the spectrum (δ = 158 (q, $^3J$ = 34 Hz) and 118 (q, 296 Hz) ppm). ESI-MS — m/z calculated for $C_{54}H_{79}N_9O_{18}$: 1141.6 u; found: $[M+H]^+$ 1142.7, $[M+Na]^+$ 1163.7.

1.4 Pre-click: [$^{225}$Ac]Ac-5B1-PEG$_8$-mcp

1) The mcp-PEG$_8$-Tz precursor was dissolved in DMSO to obtain a stock solution of $10^{-3}$ M. 1 µL (1 nmol) of the stock solution was added to 20 µL ammonium acetate buffer (0.25 M, pH=5.5) and 37 kBq $^{225}$Ac, in the form of a stock solution (0.1 M HCL), were added. The reaction mixture was placed in a thermomixer (400 rpm) at 37°C for 5 min. The radiochemical conversion was confirmed via radio iTLC.

- iTLC conditions: Glass microfiber chromatography paper impregnated with silica gel (iTLC-SG, Agilent Technologies) and aqueous EDTA (ethylenediaminetetraacetic acid, 50 mM, pH 5.5).
  Chelated $^{225}$Ac remains on the start line, free moves with solvent front.
  The iTCL was evaluated in equilibrium (+8 h). The conversion was quantitative.

2) HEPES (4-(2-hydroxyethyl)piperazine-1-ethane-sulfonic acid) buffer (5 µL, 1 M, pH 7.4) and PBS (1×, 200 µL) were added, followed by 5B1-TCO (100 µg 0.66 nmol, in 20 µL PBS). The click reaction is completed within seconds, which was confirmed via radio iTLC.

- iTLC conditions: Glass microfiber chromatography paper impregnated with silica gel (iTLC-SG, Agilent Technologies) and methanol/ acetate buffer (0.25 M, pH = 5.5) in a 60:40 ratio.
  Clicked [$^{225}$Ac]Ac-5B1-PEG$_8$-mcp remains on the start line, unreacted moves with solvent front.
  The iTCL was evaluated in equilibrium (+8 h). The conversion was quantitative.

1.5 Cells study

Before starting this assay (Supplemental Figure 4), three 6-well cell culture plates (FB012927, Fisherbrand) were seeded with $0.5\times10^6$ BxPC-3 cells per well and incubated for two days. The media was exchanged daily. On the third day, the wells were 80% confluent.

The $^{225}$Ac-labeled 5B1 antibody was mixed with cold cell media and added to two 6-well cell culture plates (1 µg antibody per well). The plates were incubated at 4°C for 3 h, followed by washing steps (3×1 mL PBS) to remove excess unbound antibodies. After 1 mL of cold media was added to each well, one plate was incubated for 3 h at 4°C, and the second at 37°C. All plates were stored at 4°C overnight before being processed.

The remaining 6-well cell culture plates were incubated with free $^{225}$Ac (1 kBq per well mixed with media) and incubated for 3 h at 37°C, respectively.

After the incubation period, the media was collected in gamma counter tubes with two wash fractions (1 mL, 50 mM EDTA, pH 7) for each well. The cells were collected in a separate tube by digesting them with NaOH solution (1 M, 2 × 1 mL). For each well, the two fractions were measured on the gamma counter directly after separation ($^{213}$Bi specific window, 440 ± 80 keV). The $^{213}$Bi release was calculated as the counts in media divided by the sum of counts (media + cells).

### 2. Supplemental figures

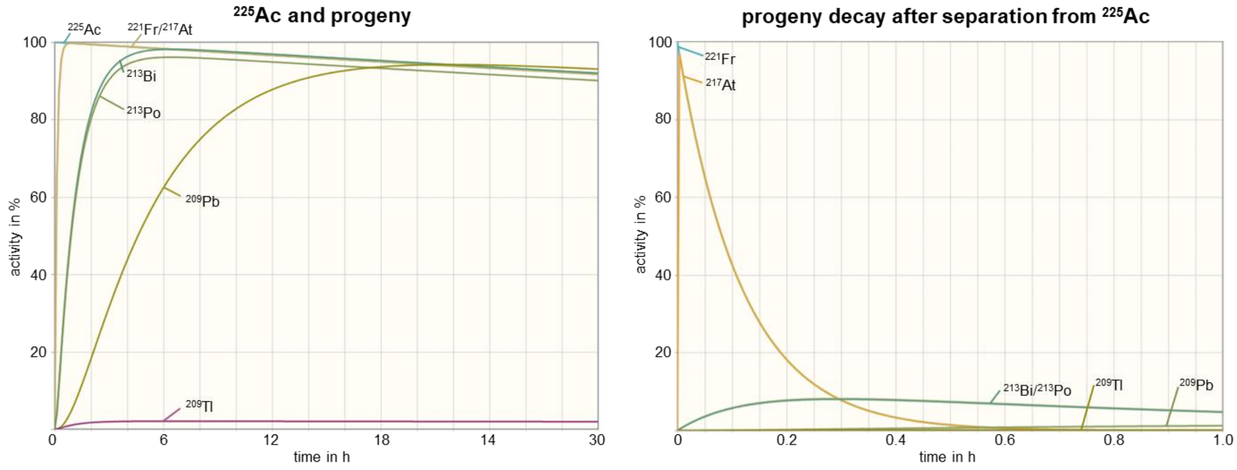

**Supplemental Figure 1.** $^{225}$Ac & progeny ingrowth.

The left graph shows the decay of $^{225}$Ac and the ingrowth of its progeny. The right graph shows the decay of $^{221}$Fr (and the ingrowth of its progeny) after separation from the mother nuclide. The plots were generated with the 'Universal Decay Calculator' (WISE Uranium Project. Accessed April 17, 2024, https://www.wise-uranium.org/rcc.html).

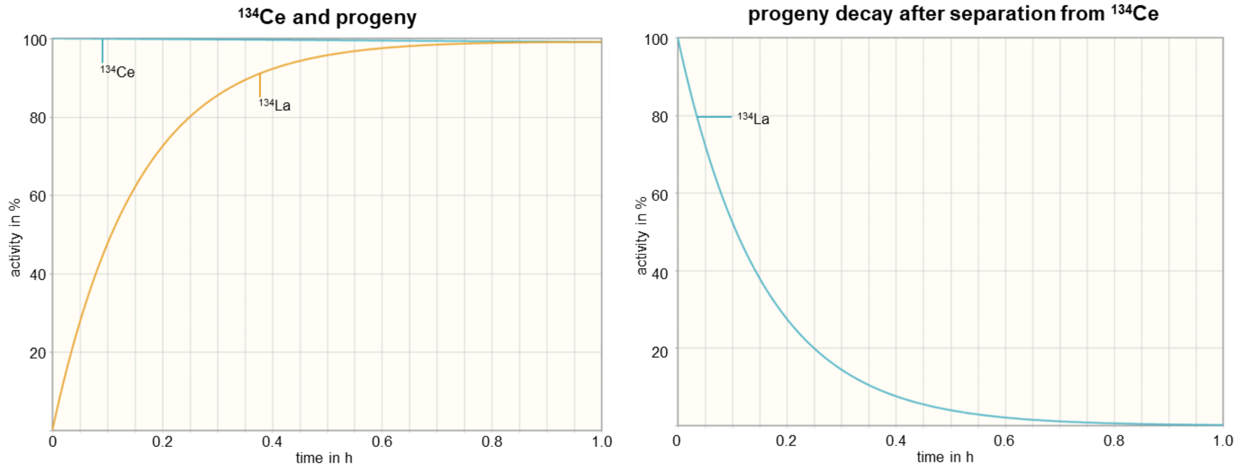

**Supplemental Figure 2.** $^{134}$Ce & progeny ingrowth.

The left graph shows the decay of $^{134}$Ce and the ingrowth of $^{134}$La. The right graph shows the decay of the progeny after separation from the mother nuclide. The plots were generated with the 'Universal Decay Calculator' (WISE Uranium Project. Accessed April 17, 2024, https://www.wise-uranium.org/rcc.html).

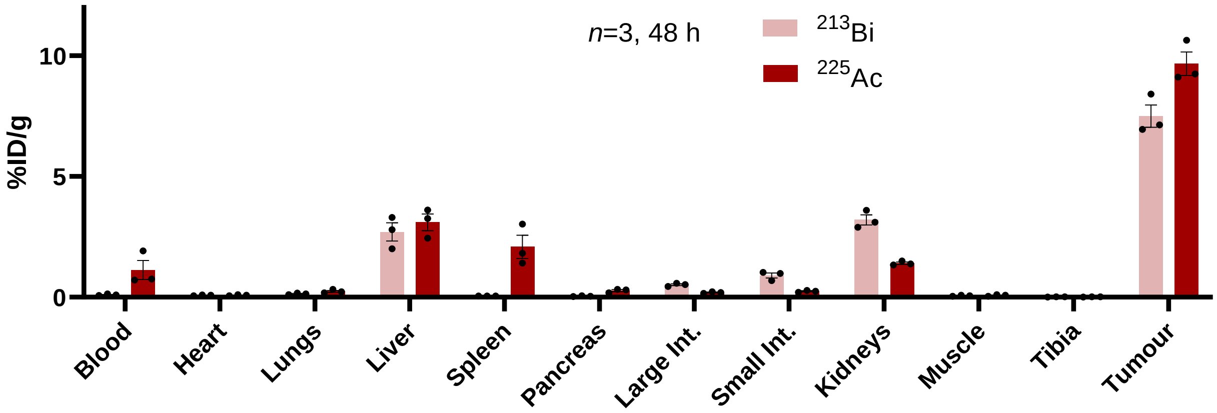

**Supplemental Figure 3.** $^{213}$Bi release in pretargeting model.

Biodistribution data of $^{225}$Ac and $^{213}$Bi at 48 h after administration of [$^{225}$Ac]Ac-mcp-PEG$_8$-Tz (2 nmol, 37 kBq) to mice (subcutaneous BxPC-3 tumours). Three days prior, the mice were pretargeted with 5B1-TCO (100 µg, 0.7 nmol). Int.=intestines.

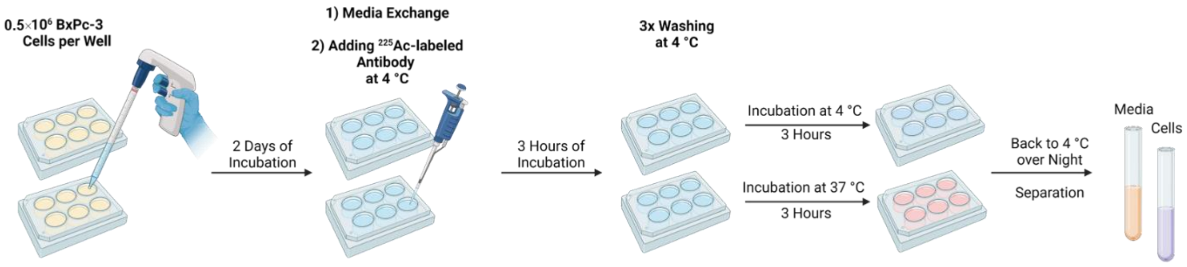

**Supplemental Figure 4** Assay design $^{213}$Bi-release study in BxPC-3 cells.

This figure was created with BioRender.

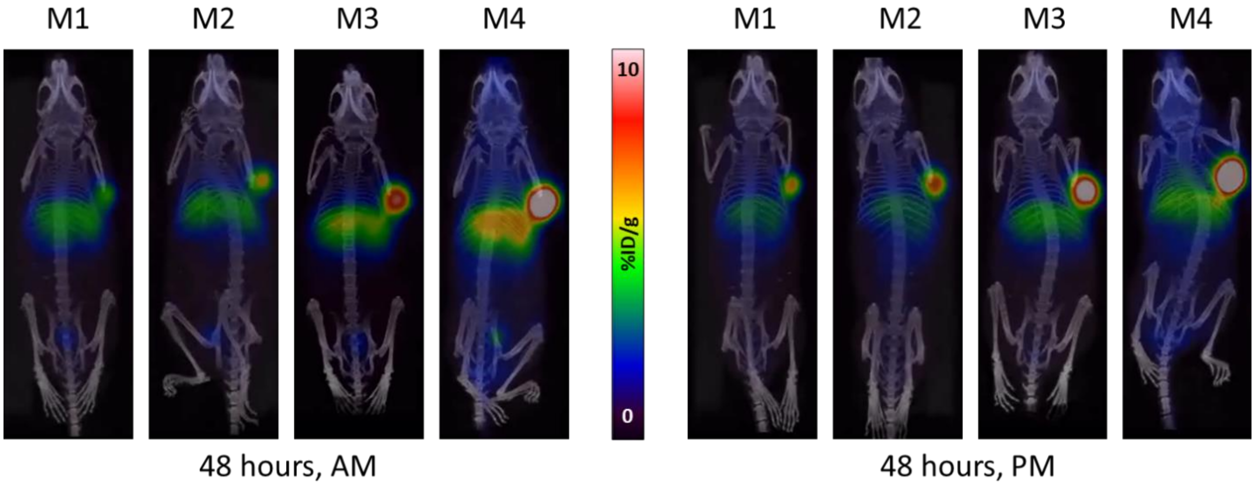

**Supplemental Figure 5.** PET images showing $^{134}$La redistribution in pretargeting study.

PET imaging (maximum-intensity projections) in four pretargeted mice receiving 3.7 MBq [$^{134}$Ce]Ce-mcp-PEG$_8$-Tz. Three days prior, the mice were pretargeted with 5B1-TCO (100 µg, 0.7 nmol).

Ante- and postmortem (AM, PM) imaging in direct comparison to visualize that, in vivo, $^{134}$La partially redistributes to the liver.

### 3. NMR spectra

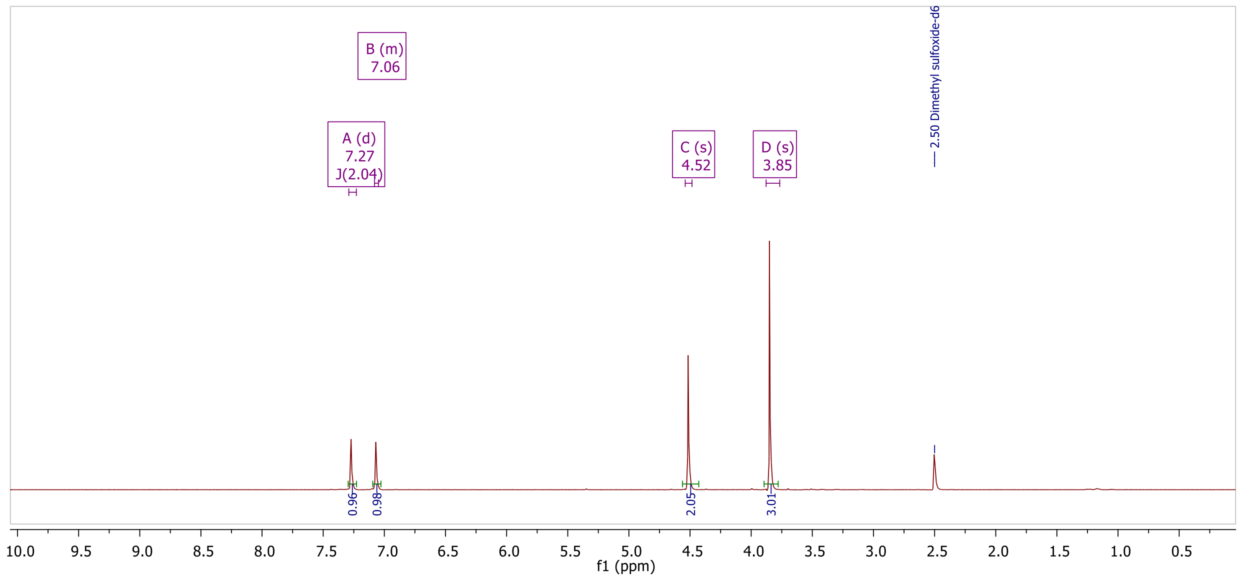

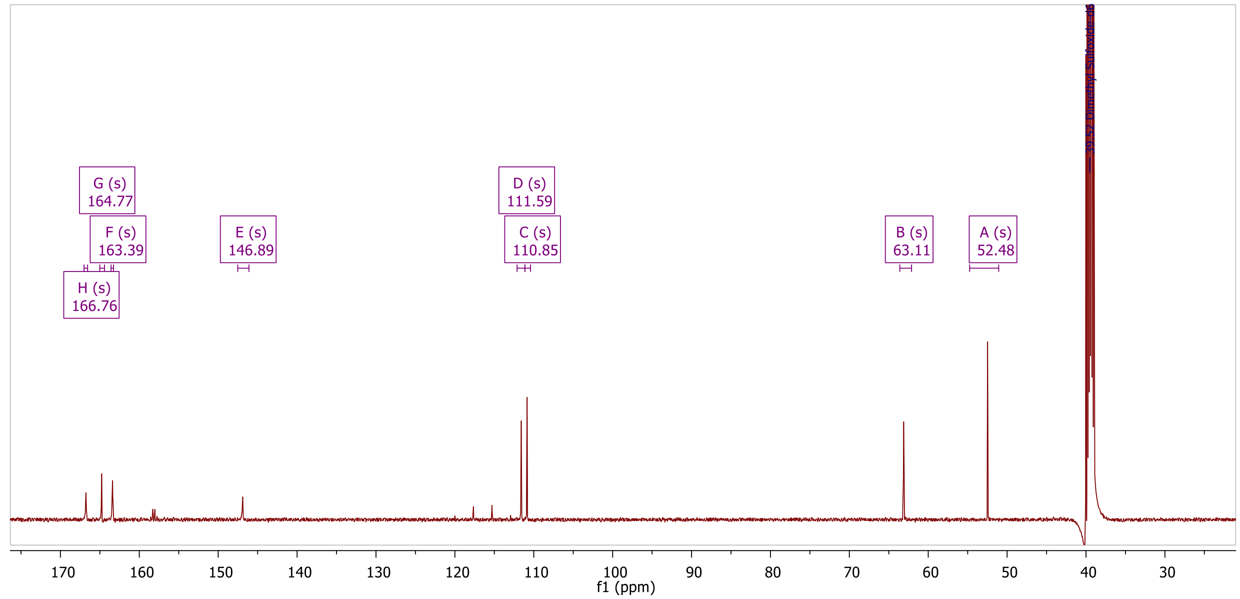

$^{1}H$ NMR and $^{13}C$ NMR spectrum of compound **C1** in DMSO-$d_6$.

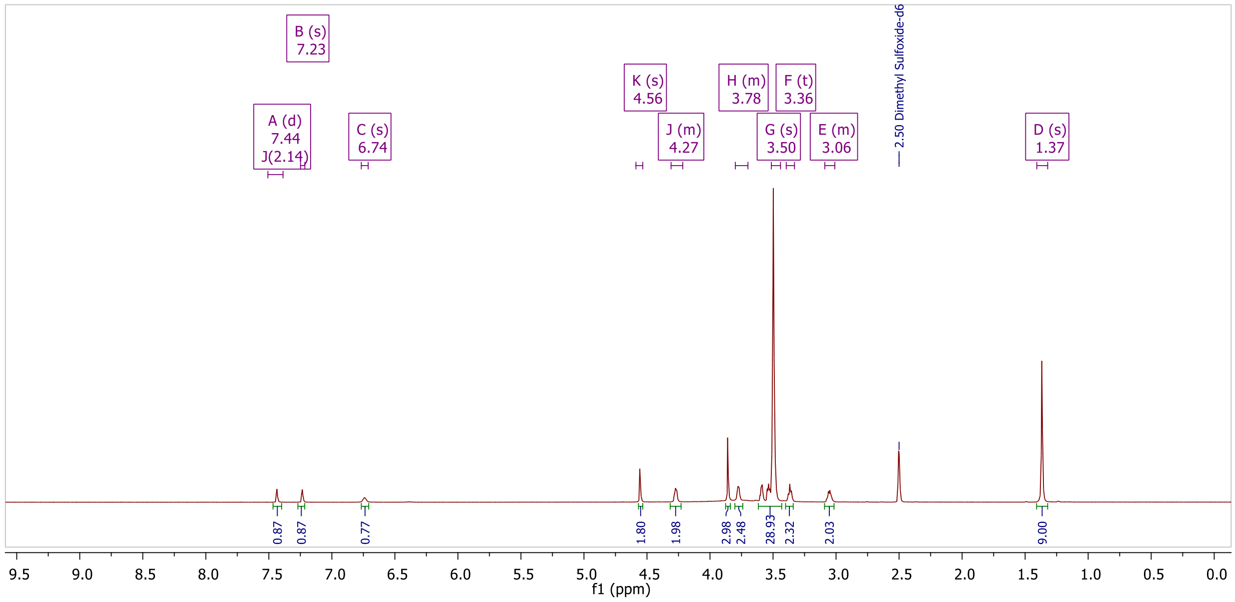

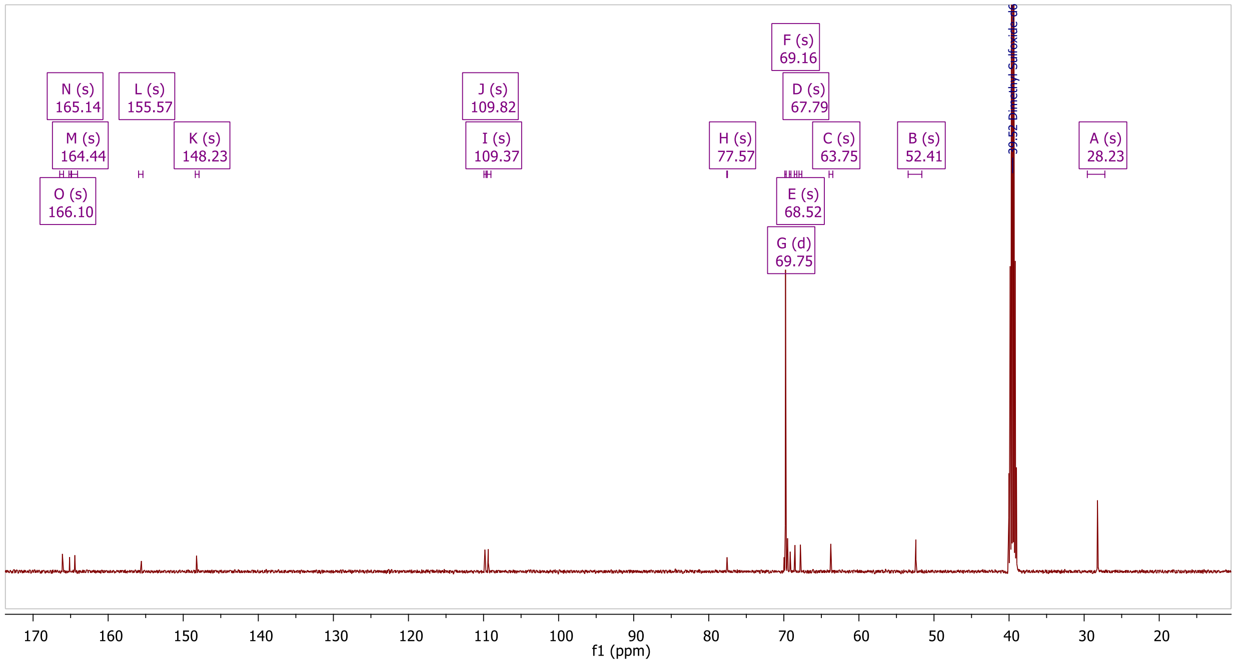

$^{1}H$ NMR and $^{13}C$ NMR spectrum of compound **C2** in DMSO-$d_6$.

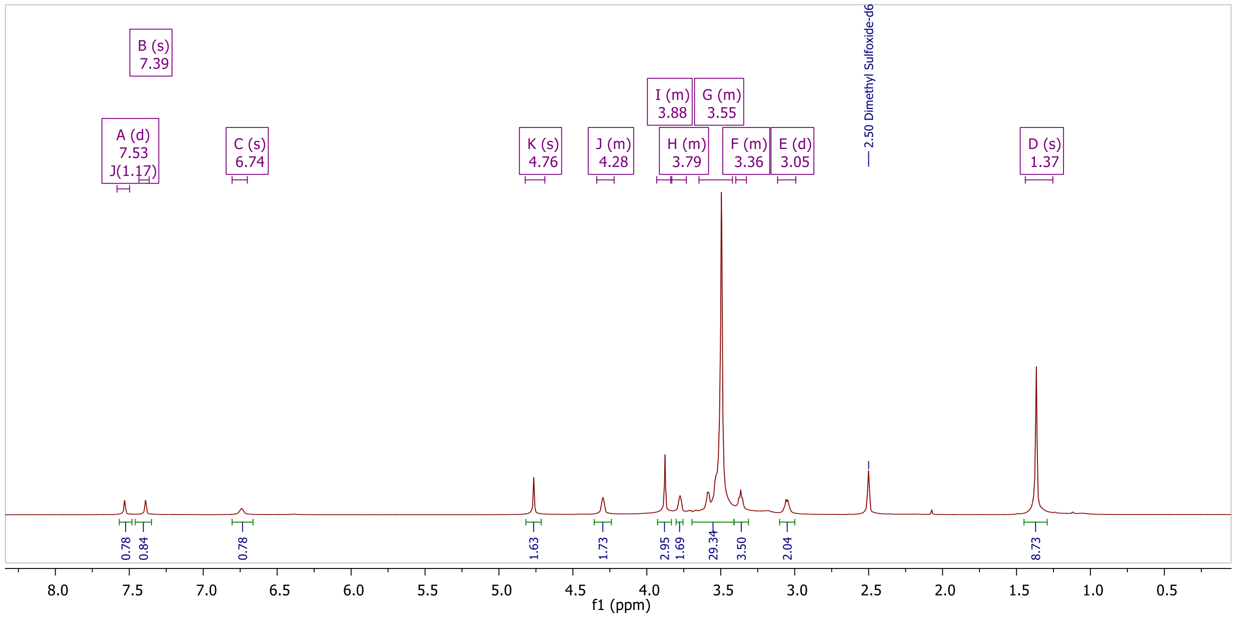

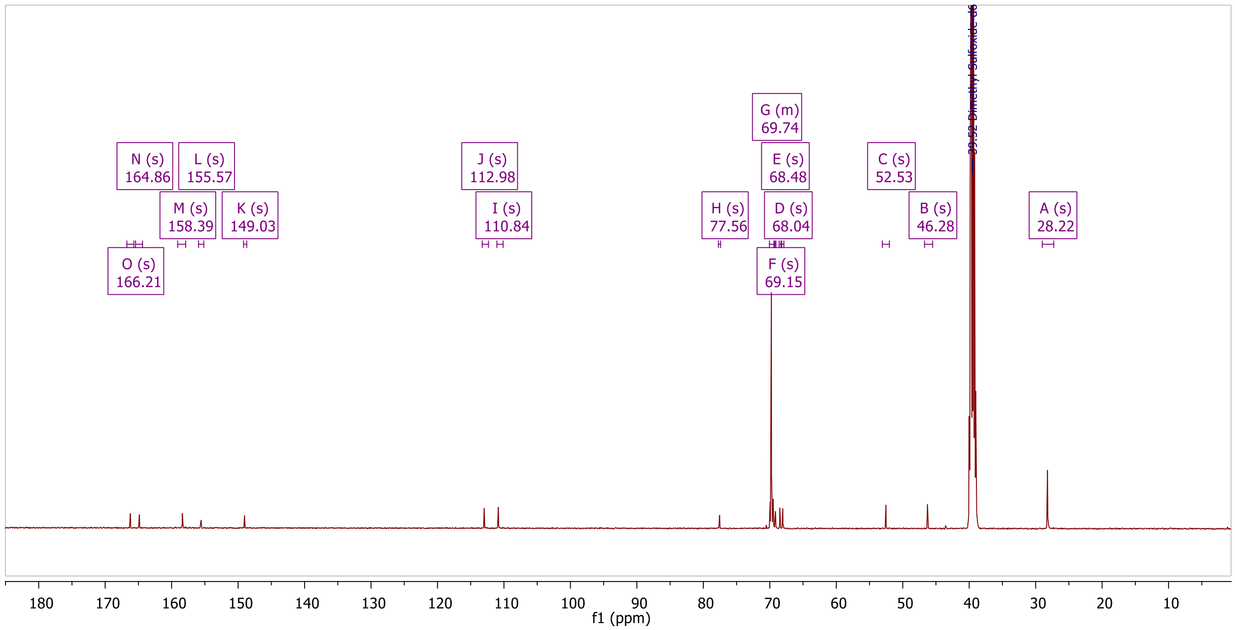

[1]H NMR and [13]C NMR spectrum of compound **C3** in DMSO-$d_6$.

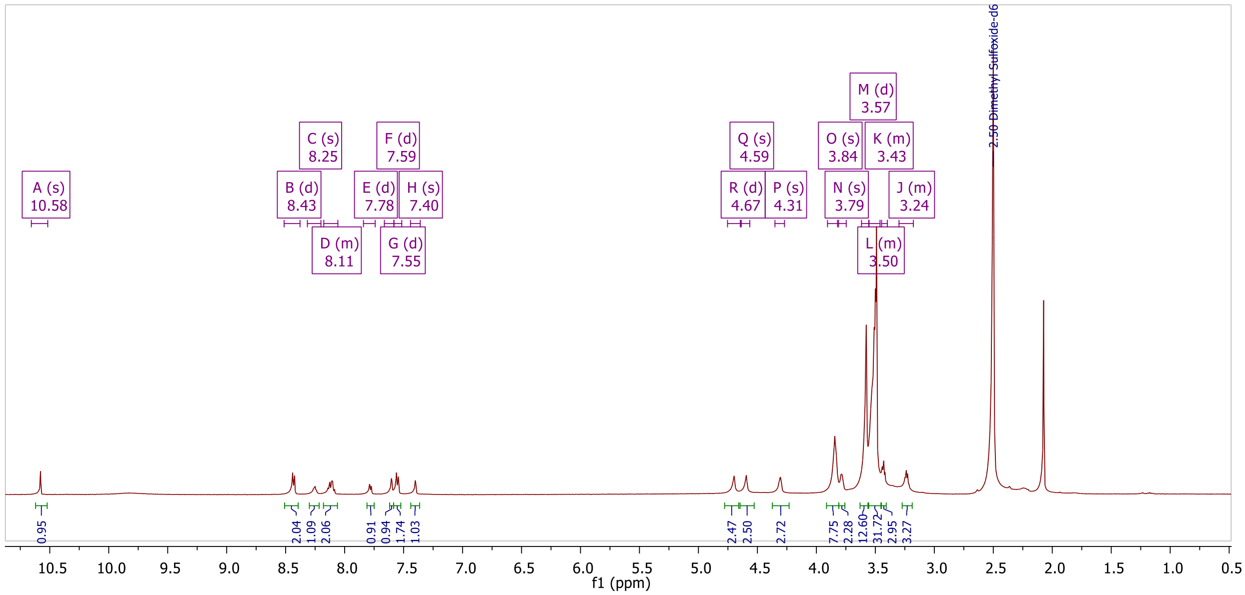

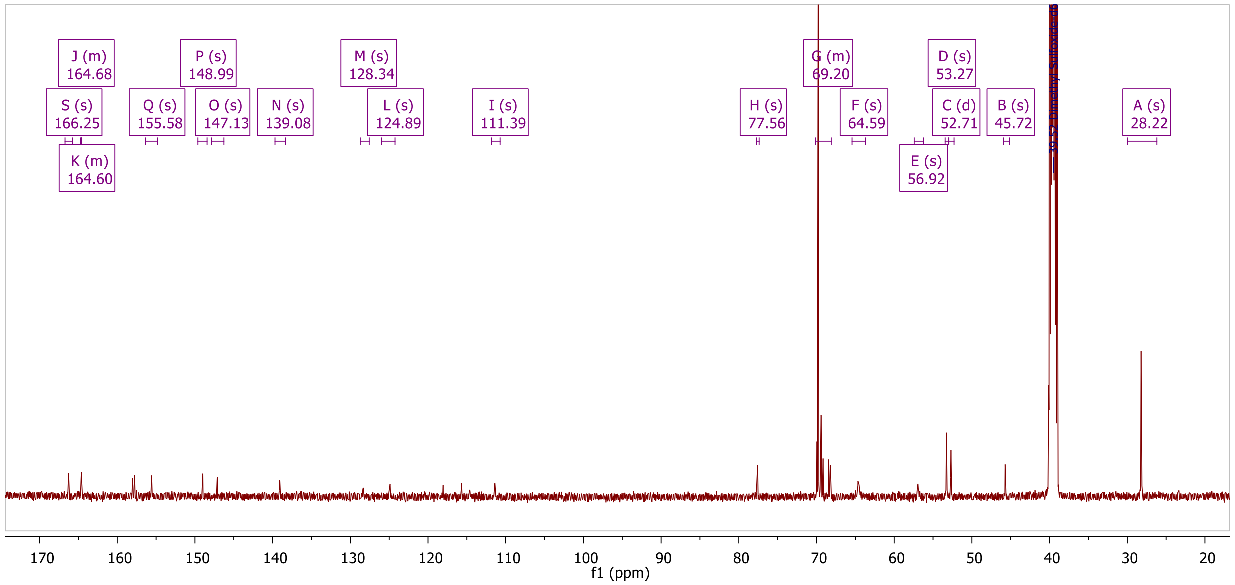

[1]H NMR and [13]C NMR spectrum of compound **C5** in DMSO-$d_6$.

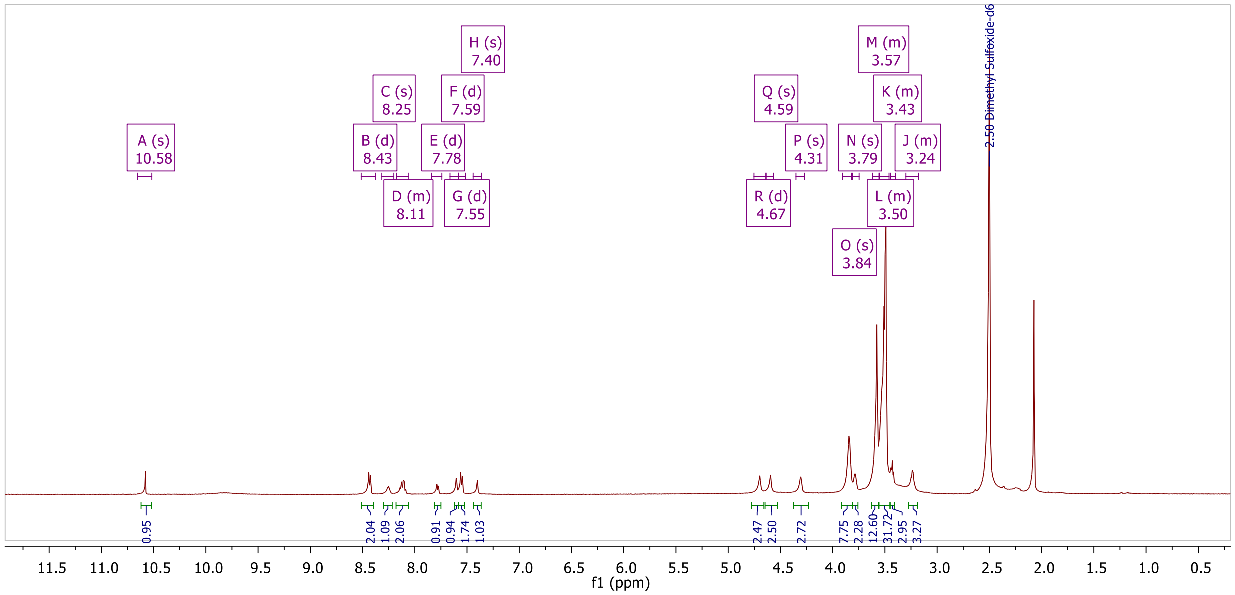

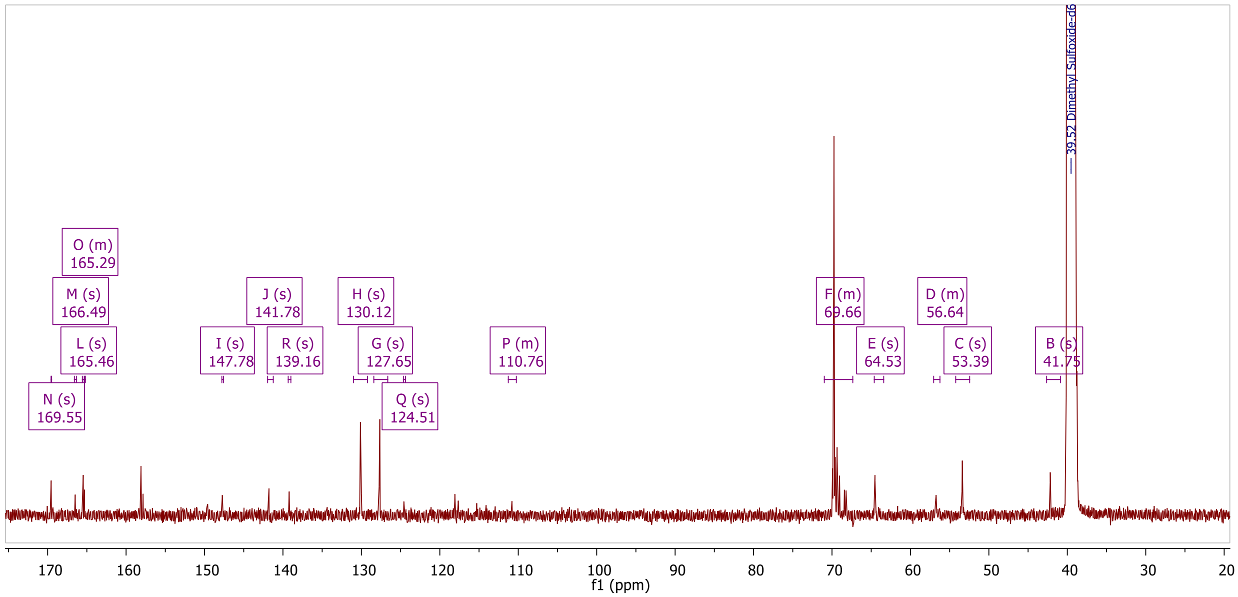

$^{1}$H NMR and $^{13}$C NMR spectrum of **mcp-PEG$_8$-Tz** in DMSO-$d_6$.
